# Believing in one’s power: a counterfactual heuristic for goal-directed control

**DOI:** 10.1101/498675

**Authors:** Valérian Chambon, Héloïse Théro, Charles Findling, Etienne Koechlin

## Abstract

Most people envision themselves as operant agents endowed with the capacity to bring about changes in the outside world. This ability to monitor one’s own causal power has long been suggested to rest upon a specific model of causal inference, i.e., a model of how our actions causally relate to their consequences. What this model is and how it may explain departures from optimal inference, e.g., illusory control and self-attribution biases, are still conjecture. To address this question, we designed a series of novel experiments requiring participants to continuously monitor their causal influence over the task environment by discriminating changes that were caused by their own actions from changes that were not. Comparing different models of choice, we found that participants’ behaviour was best explained by a model deriving the consequences of the forgone action from the current action that was taken and assuming relative divergence between both. Importantly, this model agrees with the intuitive way of construing causal power as “difference-making” in which causally efficacious actions are actions that make a difference to the world. We suggest that our model outperformed all competitors because it closely mirrors people’s belief in their causal power –a belief that is well-suited to learning action-outcome associations in controllable environments. We speculate that this belief may be part of the reason why reflecting upon one’s own causal power fundamentally differs from reasoning about external causes.

## Introduction

Inferring causality, i.e., relating changes in one variable to the causal power of another, is a general, robust, and seemingly built-in ability of the mammalian brain (Premack, 2007). The ability to draw causal inferences is critical for a wide range of behaviours and functions that range from learning and planning to flexibly adapting actions and attitudes to external contingencies (Gopnik & Schulz, 2007). Importantly, this ability may have two different types, with distinct behavioural advantages that depend on the locus of the cause itself. Thus, if the ability to draw relationships between *external* variables is paramount for adaptation and survival, it is even more so in regard to identifying *oneself* (e.g., one’s own choice or action) as the cause of a change in the world.

The ability to envision oneself as an operant agent endowed with the capacity to bring about changes in the external environment is classically referred to as “sense of agency” (Haggard & Chambon, 2012). Sense of agency builds on the biologically motivated belief that our actions are *causal* in nature: they have the power to make things happen and therefore can be implemented as an efficient means for pursuing desirable outcomes. A wealth of literature in social and cognitive psychology points towards this representation of one’s own causal power as something that is part of our natural endowment (Leotti, Iyengar, & Ochsner, 2010), develops early (Helwig, 2006) and is somewhat irrepressible (Ryan & Deci, 2006). These observations are corroborated by numerous studies showing that people readily experience control over objectively uncontrollable events (Blanco, Matute, & Vadillo, 2011), are subjected to illusions of control even when no true control exists (Langer, 1975), and experience control even though assuming control does not afford any behavioural advantage or is in fact detrimental to performance (Chambon & Haggard, 2012). The belief in one’s own causal power also comes with some advantages: higher levels of instrumental control are associated with greater general health (Bobak *et al*., 2000), fewer depressive symptoms (Rubenstein, Alloy and Abramson, 2016), and higher self-esteem (Heckhausen & Schulz, 1995). Conversely, a lowered sense of causation makes individuals more vulnerable to external, and potentially damaging, influence (Burger, 2016), and an abnormal sense of agency, such as a loss of control over one’s actions and thoughts, is long recognized as a key symptom of mental disorders (Schneider, 1959).

Questions have been raised about the function of this belief in one’s causal power. The simple fact of exercising control (i.e., of making things happen intentionally) has been suggested to be inherently rewarding (Karsh and Eitam, 2015; see also Zimbardo and Miller, 1958), as reflected by activity in a corticostriatal brain network that overlaps with the neural circuitry involved in reward and motivation processing (e.g., Tricomi, Delgado and Fiez, 2004; O’Doherty *et al*., 2004; Bjork and Hommer, 2007). Incidentally, the belief in one’s causal power is associated with an inherent *need* for control, whereby opportunities to exercise control are preferred over situations with no control, even when exercising control affords no improvement in outcome reward (e.g., Suzuki, 1999; Sharot, De Martino and Dolan, 2009; Sharot, Shiner and Dolan, 2010; Cockburn, Collins and Frank, 2014; Bown, Read and Summers, 2003; Leotti, Iyengar and Ochsner, 2010). Exercising control could serve as one of the primary means through which people foster belief in their causal power. Thus, individuals with little experience acting as an effective agent show an impaired ability to detect action-outcome contingencies, and therefore, little belief in their ability to produce desired outcomes (Leotti, Iyengar and Ochsner, 2010; Maier and Seligman, 2016; Mineka and Hendersen, 1985)^1^.

A belief in one’s causal power echoes the well-documented need in humans and animals alike to engage in activities simply to experience “competence”, that is, a sense of influencing their environment (White, 1959; see also Karsh and Eitam, 2015). Children spontaneously engage in playful exploratory behaviours where the only drive is to effect “changes “in the environment (e.g., putting a finger in a candle, knocking something off a table). Likewise, rats would readily cross an electrified grid (Nissen, 1930) and monkeys would perform costly discrimination problems (Butler, 1953) simply for the privilege of exploring and/or interacting with new territory^2^. A persistent inclination to interact with the environment has been suggested to foster action over inaction, which may prove valuable in situations in which acting does not satisfy any short-term need. A bias towards action over inaction would thus promote learning of new contingencies by favouring the acquisition of *incidental* associations between actions and action-contingent events. Once learned, these new associations could then *intentionally* be used for pursuing desirable outcomes, i.e. for achieving goal-directed behaviours (Elsner and Bernhard, 2001; see also Berlyne, 1950; Berlyne, 1966)^3^.

In addition to acquiring new action-outcome contingencies, a belief in one’s own causal efficiency may prompt the agent to probe the latent structure of the environment for causal variables. Making decisions based on the knowledge of causal variables, rather than based on local changes in the environment only, allows for better anticipation of changes in external contingencies, and for ultimately driving changes in the environment rather than being merely driven by environmental changes (Koechlin, 2014). Human cognition would be spontaneously framed in such a mode where “being a causal agent” is the default, and self-efficacy beliefs, cognitive instantiations of this default mode (Haggard & Chambon, 2012).

Collectively, the pervasiveness of this default belief in one’s causal power (Haggard & Chambon, 2012), the behavioural advantages it affords (Shapiro, Schwartz and Astin, 1996), and the various functions it underlies (Leotti *et al*., 2010) provide some clues on how human agents calculate and oversee their causal influence on the external world. A belief in the causal effectiveness of one’s action is likely to rest upon a specific mechanistic model of causal inference, i.e., a model of how actions causally relate to their consequences. The general aim of this paper is to describe what this model is. Crucially, the mechanistic model should be able to explain how people learn and update their causal influence on a trial-by-trial basis and make appropriate decisions – such as adjusting behavioural strategies to contingency changes – based on reliable causal estimates. In addition to accounting for the robustness of our everyday inferences, the model should also be simple enough to account for the *ease* with which human agents calculate action-outcome contingencies, that is, this model should be algorithmically simple. We speculate that simplicity is required to explain how control beliefs can be sustained as a default backdrop to our normal mental life (Chambon and Haggard, 2013). Finally, the mechanistic model should be endowed with properties that ultimately account for *spontaneous* illusions of control, i.e., for why people readily credit themselves for unrelated events or perceive control where there is none and act superstitiously in the belief that they are objectively controlling uncontrollable outcomes.

How people track the causal effectiveness of their actions has been a central aim of many empirical investigations, from animal learning to action cognition and personality psychology, through the prism of distinct but related, and often complementary, notions – e.g., intentional causation (Heider, 1958), perceived behavioural control (Rothbaum, Weisz and Snyder, 1982), self-efficacy (Bandura, 1989), credit assignment (Sutton and Barto, 1998), controllability (Harris, 1996), instrumental learning (Dickinson, 2001), and agency (Haggard & Chambon, 2012). Despite the great variety of disciplines concerned, three dominant approaches to instrumental causation can be distinguished, upon which relationships between action and outcome are either:

(1) retrospectively inferred (associative approach),
(2) explicitly calculated (generative approach),
(3) simply emulated (counterfactual approach)^4^

Importantly, each of these approaches draws upon different strategies with different costs and benefits. Hence, they can be distinguished on several grounds: their *efficiency*, allowing for slow or quick adaption to contingency changes; their *cost*, which makes them likely or unlikely to be implemented by resource-bounded agents; and their *vulnerability* to illusions of control and self-attribution biases. In the next section, we describe typical instances of these three approaches (associative, generative, and counterfactual models), with their respective strengths and weaknesses. Then, we turn to computational instantiations of each of these approaches, which we tested and compared across a series of modified probabilistic reversal-learning tasks. The results support the counterfactual approach motivating the development of a computational model extending the standard reinforcement-learning framework to the emulation of unseen (i.e., counterfactual) action-outcome contingencies, which allowed choices to be made online with minimal computational expense.

### Associative models: causation is about maximizing the expected value of action

One of the dominant views on causation, the associative approach, traces its roots to David Hume (1748/1978). This approach is motivated by the fact that causation is ultimately unobservable, and yet causal relations must be inferred from sensory inputs in some way (see Cheng, 1997; Walsh and Sloman, 2011; Illari, Russo and Williamson, 2011). According to Hume, only three *empirical* criteria must be met for characterizing causation: the cause must precede the outcome, the outcome must regularly follow the cause, and both must be spatially and temporally contiguous. Importantly, Hume’s definition of causation does not rely on any reference to the mechanism or process connecting events together. Causal relationships are assumed, rather than directly perceived or known, by noticing constant conjunctions between two events and by *retrospectively* presuming that a connection underpins their conjunction (Hume, 1748). Ultimately, the associative approach holds that causality is anything but a belief that is rooted in our own biological habits, that is, a pure mental construct rather than an objective property of things.

Hume’s associative account of causation has inspired various models of causal learning, from contingency models (e.g., Ward and Jenkins, 1965) to Rescorla and Wagner’s discrepancy-based learning rule (1972). A typical formulation of the associative approach can be found in studies of instrumental conditioning, in which causal action-outcome knowledge is acquired through repeated experience with event contingencies, i.e., with repeated associations between some actions (pushing a lever) and motivationally significant events, such as rewards (food delivery) (Dickinson, 2001). In the context of reinforcement learning (RL), the Rescorla-Wagner rule formalizes a simple algorithm to account for the acquisition of associative links between event representations on a trial-by-trial basis (Rescorla and Wagner, 1972; Sutton and Barto, 1998). According to this rule, association between action and consequence is learned through experiencing incremental changes in the strength of their link, and learning continues until there is no longer a discrepancy between the predicted and the actual consequences of action (Sutton and Barto, 1998). While model-free RL assesses actions sequentially through trial and error, model-based RL can include predictive knowledge (e.g., “cognitive maps”) that explicitly relate alternative actions to future environmental states (Doya *et al*., 2002). These predictive representations, which are akin to internal models of the environment, are typically learned through experiencing repeated associations between actions and effects (Daw, Niv and Dayan, 2005; Daw and Dayan, 2014). Both model-free and model-based RL algorithms operate retrospectively on experience with previous rewards by reinforcing actions that were successful in the past –i.e., by increasing the propensity to take actions that were followed by a positive reward prediction error (**Figure 1A**).

RL algorithms present several advantages that can be leveraged to model how people learn and represent their causal power. First, RL algorithms are computationally simple: they typically require only a feed-forward mapping of action to predicted consequences (Daw and Dayan, 2014). Their simplicity makes those algorithms robust and adaptive processes that can learn a variety of complex tasks even in uncertain environments (Koechlin, 2016; Gershman, 2015). This simplicity however comes at the cost of inflexibility. As an RL agent can rely only on current experience to adjust its behavioural strategies, it requires a large amount of experience to learn reliable predictions (Gershman, Markman, & Otto, 2014), and therefore, it may adapt slowly to environments exhibiting action-outcome relationships that change periodically (Koechlin, 2016).

### Generative models: causation is about inferring the latent causes generating action-outcomes links

Although causal learning exhibits many of the cardinal features of associative processes, there is evidence that human agents do not assess their causal power by simply experiencing (even repeated) conjunctions between what they want, do, and get as an action effect. Rather, they actively infer causation based on representations of latent causes generating observation. Drawing upon these internal models, agents do not only notice that effects “follow” their actions: they explicitly represent the causal sources generating action-outcome contingencies.

Various models of decision-making have endorsed this “generative” account of causation, from hidden Markov to Bayesian learning models (e.g., O’Reilly, Jbabdi, & Behrens, 2012; O’Reilly *et al*., 2013; Parr, Rees, & Friston, 2018; Mathys, Daunizeau, Friston, & Stephan, 2011; but see also Tauber, Navarro, Perfors, & Steyvers, 2017). Briefly, generative models hinge on the assumption that the observed data are the realization of one or many hidden variables or latent states (the generative source) that can be inferred with some degree of certainty, i.e., probabilistically. Crucially, generative models of instrumental causation assume that people have a more or less comprehensive representation of these hidden states, which can be learned and built up over a history of observable events, or which can be given prior to observation (e.g., O’Reilly *et al*., 2013; for non-parametric Bayesian inference, see Collins & Koechlin, 2012). Importantly, these generative representations of action-outcome relationships can be used to evaluate the different courses of action with respect to the agent’s current needs and motivational states through different causal hypotheses about action-outcome contingencies.

When inferring action-outcome relationships, there are multiple advantages for generative models. First, generative models are statistically optimal for relating observed outcomes to alternative actions as long as their representation of action-outcome mappings matches the “true” structure of the environment. Such a representation allows for a potentially optimal use of information derived from experience. Thus, rather than building on the results of the sole action taken, an agent with an accurate estimate of the outcome distribution can potentially evaluate all alternatives at once (**Figure 1B**). Second, causal relations are computed directly based on their generative representations rather than inferred based on past experience with local changes in the stimulus. The generative approach thus allows for more flexibility in adjusting to abrupt or rapid changes in contingencies as they readily occur in open-ended environments (Koechlin, 2016).

Shortcomings of the generative approach concern both its computational cost and its biological plausibility. Under a Bayesian setting, the generative approach assumes that the agent can build an exhaustive generative model of all possible states on which the inference is drawn. However, in real-life situations, representing and updating all possible alternatives at once leads to intractable computational costs. This makes the *complete* generative solution unlikely to be implemented by the brain (Eckstein *et al*., 2004), which explains why people often depart from statistically optimal predictions made by normative models (e.g., Waldmann and Walker, 2005; see also Blanco, Matute and Vadillo, 2011; Gershman, 2015). Interestingly, departures from normative predictions often arise in the form of illusions of control in which people behave superstitiously in the belief that they are controlling uncontrollable outcomes (Langer, 1975), such as those occurring when contrasting instrumental vs. observational learning (Waldmann & Hagmayer, 2005) and naturalistic vs. analytic contexts (Matute, 1996), or when experiencing imposed vs. chosen gambling outcomes (Kool, Gatez, & Botvinick, 2013). Generative models have difficulty accounting for such illusions while at the same time failing to address causal problems that human subjects easily solve (Sloman & Lagnado, 2015). Therefore, questions have been raised about whether deviations from normativity only capture *approximations* of the true generative solution – e.g., due to limits on the size of working memory or on the quantity of attentional resources – or whether they ask for a rethink of how individuals construe their causal power, i.e., with relatively high efficiency and sustainable computational costs (e.g., Jones and Love, 2011; Markman and Otto, 2011; Bowers and Davis, 2012; Collins & Koechlin, 2012).

**Figure 1.**
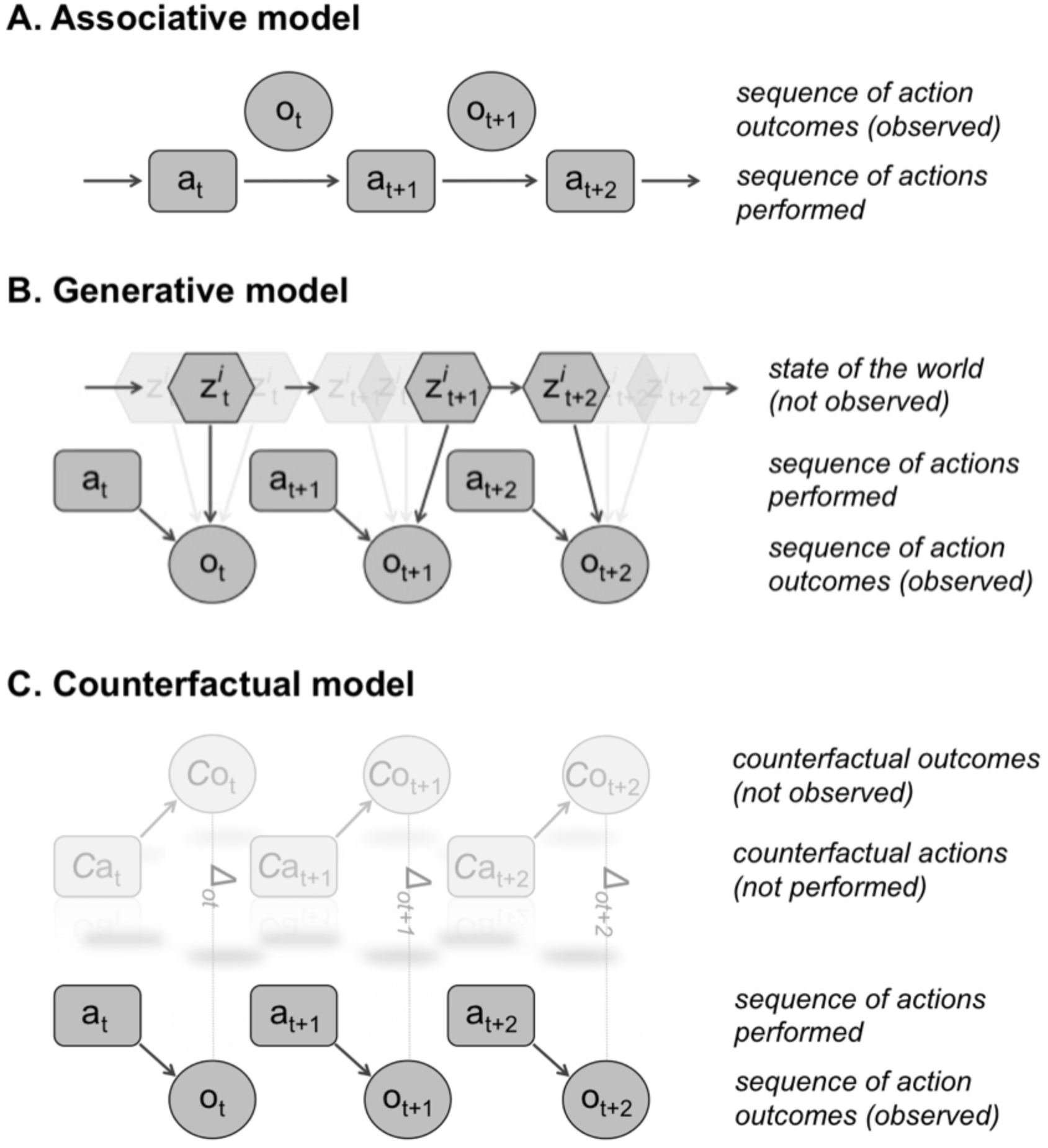
Three decision-making and learning models to account for how human subjects infer and monitor their causal power. **A. *Associative model:*** agents associate action (A) and outcome (O) through experiencing repeated event contingencies, and reinforce actions that were successful in the past. Agents learn preferences for actions without ever explicitly learning or reasoning about the (hidden) structure of the environment. **B. *Generative model:*** agents infer action-outcome causal relationships based on an internal “model” of the world (Z; the “generative” source) that explicitly relates actions (A) to future outcomes (O). Generative models can ideally learn all possible hidden states (octagons in transparency) relating the action performed with the observed outcome. **C. *Counterfactual model:*** agents simulate what would have happened (*C*_o_) had another action (*C*_a_) been taken. Under the counterfactual view, an action has causal power over an observed outcome if a change in that action (i.e., another action, or no action, is taken) leads a change in the outcome. Ideally, causal actions are those maximising the difference (Δ_o_) between factual and counterfactual outcomes.

### Counterfactual models: causation is about actions that make a difference

Associative algorithms describe agents that can adapt to the causal structure of the world with minimal computational expense while generative models directly infer causation by relying on explicit representations of latent causes generating action-outcome relationships. Hence, associative and generative models of causation stand as opposite extremes on a continuum between statistical efficiency and computational tractability. Importantly, a number of theoretical and empirical works have suggested that counterfactual reasoning might sit in the middle of this continuum.

In the decision-making domain, *counterfactual reasoning* (CF) draws upon representations of what would have happened had another choice been made (e.g. Boorman, Behrens and Rushworth, 2011). If a large psychological literature has chronicled the affective consequences of counterfactuals, especially regret, on choice behaviour (Bell, 1982; Roese, 1997; Coricelli *et al*., 2005), there is also abundant empirical evidence that people generate counterfactuals, i.e., simulate alternative possible events and their outcomes, when they think about causal relations. Thus, when a change to an event leads to a change in the outcome, people rate it as more causal than when a change to the event would not undo the outcome (e.g., Walsh and Sloman, 2011). Similarly, making a counterfactual alternative available strongly influences causal judgements, so that the greater the number of counterfactual alternatives for an event, the more causal this event is perceived (Spellman and Kincannon, 2001; McCloy and Byrne, 2002; Byrne, 2005). While CF plays a role in causal reasoning, it is, however, not equally applied to all types of situations. People are more prone to counterfactual thinking for causal relations that have a behavioural significance to them, such as voluntary actions (see Roese, 1997, for a review). Thus, individuals are more likely to generate counterfactuals when judging causation in situations involving *actions* than inactions (“agency effect”, see Byrne, 2002) as well as *controllable* events (e.g., voluntary choices) instead of uncontrollable events (e.g., an asthma attack) (Girotto, Legrenzi and Rizzo, 1991; N’Gbala and Branscombe, 1995). Conversely, decreasing causal power and personal control diminishes the propensity for counterfactual thinking (Scholl and Sassenberg, 2014). Together, these results suggest that there is a close relationship between counterfactual thinking and people’s sense of causation for actions under their direct control.

Importantly, the CF account defines a cause as something that makes a *difference* to another event (i.e., the outcome would have been different had another action been performed), which endorses a very intuitive way of construing causation as *difference-making*. In the counterfactual literature, models of causal reasoning (e.g., Pearl, 2000) share this idea with modern instantiations of the associative approach, such as recent accounts based on experienced action-outcome contingency – where contingency is defined as the difference between conditional probabilities, such as the so-called “ΔΡ rule” (e.g., Tanaka, Balleine and O’Doherty, 2008) – and with model-based learning algorithms drawing upon the notion of instrumental divergence (i.e., “Jensen-Shannon divergence”). Instrumental divergence formalizes the causal power of an action as the difference between probabilities of a given outcome in the presence vs. absence of this action (Liljeholm *et al*., 2011; Liljeholm *et al*., 2013; Mistry and Liljeholm, 2016). Interestingly, both counterfactual reasoning and instrumental divergence are endowed with the same prior belief about goal-directed actions. They assume that goal-directed actions are instrumental in nature: choosing action A over action B (or choosing to act vs. not acting) *makes a difference in terms of the outcome*. Additionally, the greater the action differs with respect to its contingent states (the factual and counterfactual outcomes), the more flexible control the subject has over the environment (**Figure 1C**).

Importantly, both CF reasoning and instrumental divergence imply being able to *emulate*^5^ the outcome associated with the unchosen course of action, i.e. to adapt behaviour *as if the unchosen course of action had been carried out and its outcome had actually occurred*. Both views make the same assumption: when the difference between the *real* and *emulated* outcome is maximal, instrumental control (hence, causal power of one’s action) should be maximally assumed. Counterfactual emulation offers several advantages over both the associative and generative approaches. For example, CF makes it possible to learn information from *unchosen* alternatives without having to incur the costs of taking the alternative course of action (Boorman, Behrens and Rushworth, 2011; Buchsbaum *et al*., 2012; Lohrenz *et al*., 2007; Collins & Koechlin, 2012). Counterfactual emulation is also far less costly than statistical inferences within generative models based on multiple hidden causes. Unlike generative models that must *learn* causation through considering and updating all possible alternative causes at once, CF assumes causation through a simple prior belief based on difference-making.

## Overview of the present study

CF studies have provided convincing evidence that people generate counterfactuals when reasoning about causation (Sloman and Lagnado, 2015, for a review), whereas instrumental divergence provides a learning rule for how people make choices based on *maximized* divergence (Mistry & Liljeholm, 2016). However, both views have shortcomings. So far, CF models of causal reasoning have only been applied to static environments and abstract settings –i.e., verbal scenarios or summary descriptions of causal situations–, while studies drawing upon instrumental divergence critically lack of an algorithmic insight into *how* unchosen situations are emulated and according to which rule (e.g., what value should be assigned to the alternative? How can this value be learned and according to what dynamics? And what should be its update rule?). In this paper, we propose to bridge the gap between these two approaches by building and testing a counterfactual model addressing these issues.

We tested and compared the performance of this model (hereafter, CF) against various instantiations of the associative and generative classes (hereafter, RL, BM, BC) in a series of tasks in which there was some uncertainty about the identity of the causal agent (**Figure 2A**). The tasks were built on a modified reversal-learning procedure (Rolls, 2000) and modelled a dynamic environment where action feedbacks were intrinsically noisy and instrumental or environmental contingencies could change unexpectedly (**Figure 2B**). To maximize their performance, subjects had to continuously monitor their causal influence over the task environment, by discriminating changes that were caused *by their own actions* from changes *that were not*. In turn, discriminating self-from externally caused outcomes required tracking changes in the different statistics manipulated in the task (i.e., action-outcome dependency, value, variance) and to flexibly adjust to these changes.

## Overview of the experimental paradigm

We ran three distinct experiments in three different groups of participants. In each experiment, the task consisted of a *two* slot-machines game (**Figure 2A**). Thus, for each trial, the participants had to make two distinct choices: (i) first, selecting which of the two machines she wanted to know the result of, then (ii) selecting which of the two buttons (a square or a circle) to press in order to trigger the machines. The order of choice (see **Figure 2A**: machine then button, or button then machine) was counterbalanced within participants, while the spatial mapping of the task stimuli and the response keys were counterbalanced across participants.

Crucially, the participant was informed that although she played the two machines simultaneously, she would control *one and only one machine*. Thus, for one machine only, whatever the button pressed (a square or a circle) the average reward was the same, whereas for the other machine, one button (the “best-rewarding” button) gave a higher reward on average than the other (the “least-rewarding” button). Put another way, the chosen button influenced the gains of one machine only (the “controlled” machine), whereas the gains of the other machine (the “non-controlled” machine) were independent of the button pressed by the subject. To maximize her final payoff, the participant had to determine which machine she controlled, that is, the machine for which there was a best- and a least-rewarding button.

The participant was informed that she would always win the sum of the gains from *both* machines on each trial. This was to motivate her to track the controlled machine (i.e., the machine for which her choice made a difference) rather than systematically searching for the best-rewarding machine. After a given number of trials, a feedback screen displayed her current payoff, which was graphically represented as the sum of the gains produced by each machine during these last trials.

Gains produced by each bandit machine were drawn from Gaussian probability distributions (truncated between 1 and 100 and then rounded to the nearest integer). Mean, variance, and “divergence” of these distributions varied across conditions. Divergence refers to the distance between gains distributions associated with each button or machine. The divergence constituted our measure of control. The machine with a *positive* divergence was the *controlled* machine, that is, the machine for which there was a (maximal) difference in the probability distribution of gains associated with each action (e.g., **Figure 4A**, red and green distributions). Conversely, the *non-controlled* machine was the machine with a *null-divergence;* that is, the machine for which each action was similar with respect to its contingent state (e.g., **Figure 4A**, grey distribution). Thus, instrumental divergence defines the ‘controlled’ machine as the machine for which making a choice (e.g., selecting button A vs. B) makes a difference in terms of the outcome, in accordance with various accounts of instrumental causation as “difference-making” (e.g., Walsh and Sloman, 2011; Liljeholm *et al*., 2013; Beebee, Hitchcock and Price, 2017). It is worth noting that instrumental divergence quantifies the degree to which alternative actions differ with respect to contingent states, and hence this approach is formally equivalent to another highly related information theoretic measure, mutual information, which quantifies the statistical dependency between an action and a subsequent event (see Liljeholm *et al*., 2013).

Finally, participants were informed that unpredictable reversals could occur during the task so that either buttons or machines reversed unpredictably from time to time (e.g., the best-rewarding button became the least-rewarding button, or the controlled machine became the non-controlled machine) (**Figure 2B)**. Participants were thus explicitly asked to pay attention to the relationship between their gains and their choices so they could identify these reversals as fast as possible and adapt their choices accordingly.

Importantly, the experimental conditions differed in how these reversals were implemented. Thus, depending on the condition within each experiment, participants had to monitor reversals in either:

(i) the **statistical dependency** between their action and the resulting outcome, or (ii) the **rewarding value** of the outcomes produced by each machine, or (iii) the **variability** of these outcomes over time.

The first experiment tested the influence of these statistics on the participant’s choice separately, i.e., within independent experimental sessions. In this experiment, either *explicit* (Expt. 1a) or *implicit* (Expt. 1b) instructions were given to participants about their actual control over the task. The second experiment (Expt. 2) implemented the same procedure but controlled for *interaction effects* between the 3 statistics that were manipulated (e.g., dependency, value, variability) by employing a full factorial design in which these statistics were systematically crossed.

**Figure 2.**
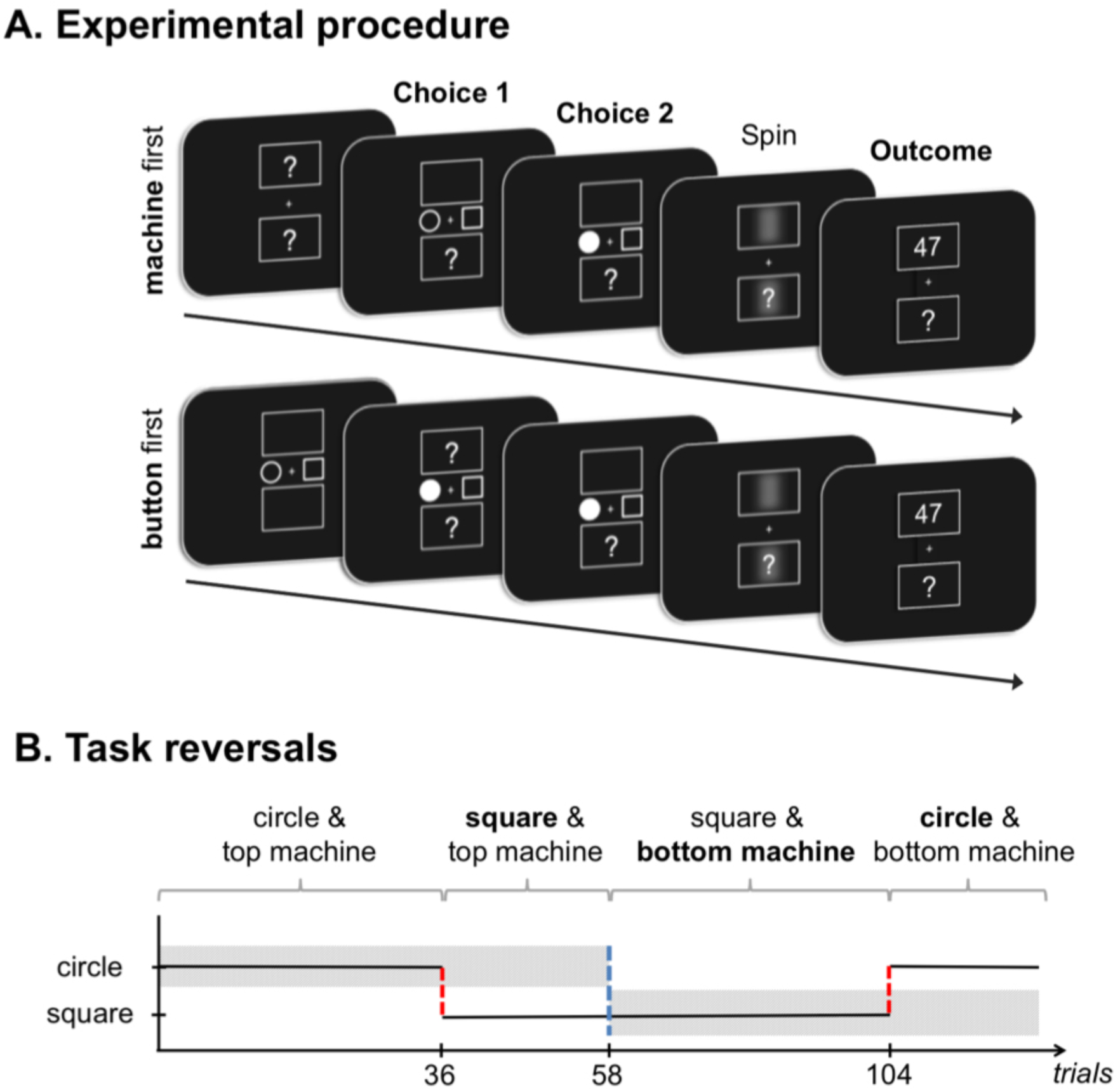
Schematic of trial procedure and stimuli. **(A)** A trial started with the presentation of two bandit machines above and below a central fixation. In one half of the blocks, the subject had to first select a machine (here, the top machine, **Choice 1**, top panel) and then a button (here, the left button, **Choice 2**, top panel), and conversely in the other half (button, then machine; see bottom panel). Note that only the gains of the selected machine were displayed at the end of the trial. Each trial lasted approximately 3s. **(B) Schematic of reversals during the task**. The solid line represents the best button during the ongoing block, whereas the grey rectangles represent the location of either (i) the controlled machine (expt. 1: dependency session), (ii) the best-rewarding machine (expt. 1: value session), or (iii) the low-variable machine (expt. 1: variance session). The vertical red dashed lines signal a reversal on the best-rewarding button (circle to square, or the converse) whereas the vertical blue dashed line signals a machine reversal. In all experiments, “button” or “machine” reversals occurred after a variable number of trials.

## Modelling

In both experiments, four classes of models were built and fitted to participant’s choices: *(i)* a simple reinforcement learning model (RL), *(ii)* a counterfactual learning model (CF) built in a model-free reinforcement learning framework, and two generative models whose aim was to learn the task environment *correctly* by searching for either *(iii)* the “best-rewarding” state (Bayesian-maximizer, BM) or *(iv)* the “controlled” state (Bayesian-controller, BC) in the task environment. Each of these four models draws on different assumptions about how subjects’ beliefs are formed and updated on a trial-by-trial basis and therefore makes different predictions on how choices are made based on these beliefs.

In all four models, the *two-stage* decision process was concatenated into one single decision made between the four possible combinations of machines and buttons. We did so because reaction times suggested that the two successive choices (machine then button, or button then machine) were chunked into one unique choice made between four action sequences. Indeed, in all tasks, the reaction times for choices were significantly slower for the first choice made, whether this choice was a button (paired t-tests, all experiments: all *t*’s(15–25) < −2.91, all *p*’s < 0.007) or a machine (all experiments: all *t*’s(15–25) < −11.1, all *p*’s < 0.001). Under these conditions, it has been shown that modelling two successive choices as one unique decision better predicts the participants’ data (Solway and Botvinick, 2015; Dezfouli and Balleine, 2013).

In all four models, each of the 4 possible actions made by the participant on each trial (choosing between 2 buttons × 2 machines) was associated with either an *action value* for both RL and CF models (**Figure 3**), or with *beliefs* (indexing the probability to be in one particular state among all possible states) for the generative models (**Figure 4**). All four models went through the same two steps on each trial. The first step consisted of updating the internal value or the beliefs associated with each of the 4 possible actions, depending on the outcome obtained in the previous trial. The updating rule was different between models (see below). The resulting internal values or beliefs were then used to compute the probability to choose one action over its 3 alternatives. The second step consisted in making a choice based on either internal values or beliefs using a non-deterministic (*softmax*) decision rule (see below, “**Action selection**”).

### RL model

Each of the 4 possible actions were associated with an internal value (Sutton and Barto, 1998), which is also called an *action-value* (**Figure 3**, top panel). The values themselves are hidden but are thought to drive choices between alternative actions. Specifically, the model draws upon the notion of prediction error (δ), which measures the discrepancy between actual outcome value, called *reward* (R) here, and the expected outcome for the chosen action (i.e., the chosen *value*) at time step *t*:

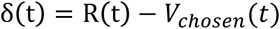

According to the Rescorla and Wagner’s rule (1972), such a prediction error is used to update the value of the chosen action, as follows:

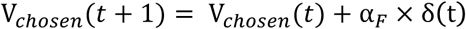

α_F_ is a fitted parameter capturing the rate at which prediction error updates the action values, and thus it is called the (factual) learning rate. Action values represent the reward value expected for choosing this particular action. Here, the action values associated with the three *unchosen* actions are kept constant (i.e., they are not updated):

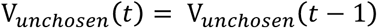

### CF model

Contrary to typical RL, values of the *unchosen* actions (i.e., counterfactuals) were explicitly updated in the CF model, and this update was performed according to a specific dynamic (e.g., learning rate). Note that counterfactual rewards were not experienced or seen, and therefore they must be somehow inferred by the participant. CF models assume that such inference requires emulating the unchosen action *as if* it was effectively taken and to derive the corresponding counterfactual outcome from it (**Figure 3**, bottom panel). Some uncertainty remains about how to model the emulation process. Converging evidence from reinforcement comparison methods (Sutton, 1984; Dayan, 1991; Kaelbling, Littman, & Moore, 1996) and behavioural economics (Palminteri *et al*., 2015; Denrell, 2015; Burke *et al*., 2016) suggests that people always make decisions relative to a context-dependent reference. When the context is a (binary or continuous) distribution of gain and losses, this reference approximates the mean of the distribution (Palminteri *et al*., 2015; Kahneman and Miller, 1986). Interestingly the mean is an important, often optimal, operator that allows for minimizing prediction error in error-prone situations, i.e., under uncertainty (de Gardelle and Summerfield, 2012). In the following, counterfactual rewards were thus inferred based on a simple contextual rule. The counterfactual reward (*R_CF_*) was derived from the actual reward, which it mirrored through a *reference* point (*P*) approximating the mean of the underlying generative distribution. The value of this reference (or “context value”, Palminteri *et al*., 2015) was separately fitted, rather than fixed or learned from reward history, in each participant:

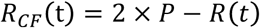

According to this rule, when participants obtained a high reward (“high” being defined as being above the reference reward), the counterfactual reward associated with the unchosen action was inferred as being a “low” reward (i.e., below the reference reward), and the probability to *stay* with the same action on next trial increased. Conversely, when the obtained reward was low, the counterfactual reward was inferred as being “high”, and the probability to *switch* action on next trial increased. The emulated counterfactual reward thus allowed for computing a *counterfactual prediction error* (δ*_CF_*) and a *counterfactual learning rate* (α*_CF_*), which was used to update the value of the unchosen actions according to a generalized version of the Rescorla & Wagner’s rule, as follows:

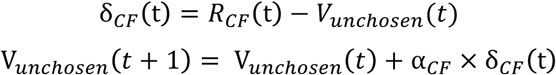

Note that because participants chose between *four* possible actions, there were necessarily *three* unchosen actions for each choice made: (i) the unchosen button associated with the chosen machine, (ii) the chosen button associated with the unchosen machine, and (iii) the unchosen button associated with the unchosen machine. Hence, the model was endowed with 3 counterfactual learning rates (α*_CF_*_1_, α*_CF_*_2_ and α*_CF_*_3_), which were fitted to each participant separately.

**Figure 3.**
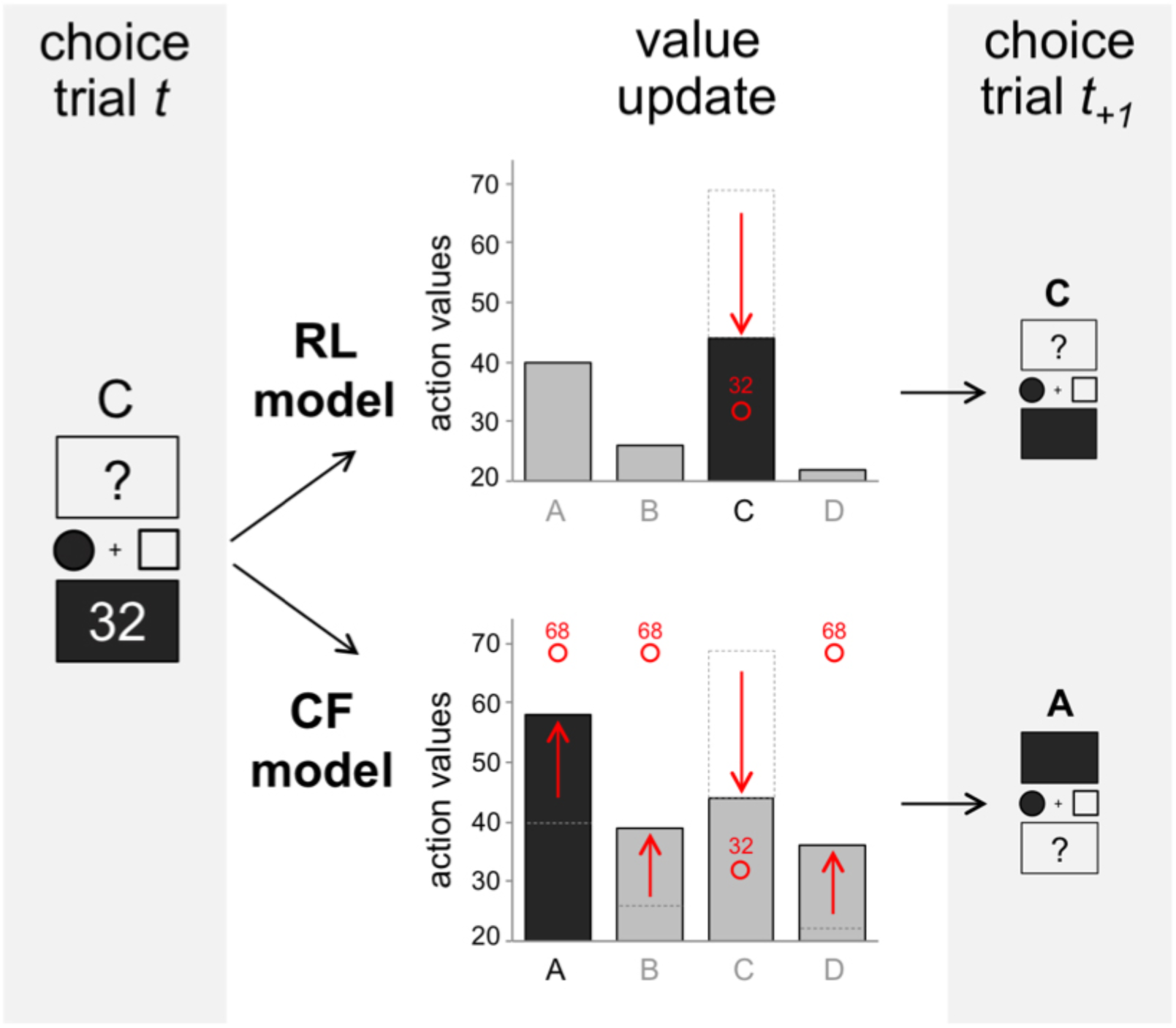
Schematic of the two-stages decision process in RL and CF models. On trial *t*, the circle button and the bottom machine are chosen, and ‘32’ is obtained as a reward. RL (top) and CF (bottom) models differ in how action values are updated. While both models use the current reward to update the value of the chosen action through the Rescorla-Wagner (R-W) rule, only the CF model updates the value of the *unchosen* actions. The CF model derives a fictive counterfactual outcome (here, ‘68’) from the actual outcome (‘32’), which it mirrors through a reference point approximating the mean of the underlying reward distribution. The counterfactual outcome is then used to update the value of the unchosen actions through the classical R-W rule. Importantly, these different updates can lead the two models to make different choices: on the next trial, the RL model chooses the circle button and the bottom machine (choice **C**), while the CF model chooses the circle button and the top machine (choice **A**). Note that the figure shows a reversal in contingency (C is no longer the best valued action). As can be seen, the CF model adapts quickly to the reversal (it now chooses **A**), whereas the RL model sticks to the same action (it keeps choosing **C** as before).

### Generative model

The generative model was a Bayesian learner that updated beliefs, and not values, associated with each possible action, on each trial. Here, a belief referred to a *probability* for an action to be in a given state (**Figure 4**, and **Appendix A**, “Generative model”). Instructions that were explicitly given to participants defined four possible states associated with three generative distributions (G):

1. the state associated with having selected the *best-rewarding* button of the *controlled* machine (*G_1_*),
2. the state associated with having selected the *least-rewarding* button of the controlled machine (G*_2_*),
3. the state associated with having selected the non-controlled machine (*G*_3_).

On each trial, the model aimed to infer the correct state/action pair, i.e., to infer which among the three possible distributions generated the observed outcome given the button pressed. The model then updated its belief about all state/action pairs along with the parameters (mean, standard-deviation) of each generative distribution, given the new observations. Our model was implemented with a specific task structure defining the number of possible states (the three generative distributions), actions (the four possible actions), and hidden variables to describe them (e.g., the mean and standard-deviation of the generative distributions). The model assumed the generative distributions to be Gaussian with fixed mean and standard deviation. On each trial, the mean and variance of each generative distribution was inferred by the model, which was based on the history of observations, through Bayesian inference (see **Appendix A**, “Generative model”, for details). As reversals between actions occurred, the model also needed to infer a volatility parameter, the volatility being the probability for the states to reverse between actions. Thus, for each trial, the Bayesian models needed to infer a set of 7 parameters: the three Gaussian means, the three Gaussian variances, and the volatility parameter (**Figure S1**).

To test how participants interpreted our instructions, we built two different Bayesian models: a Bayesian-controller (BC), and a Bayesian-maximizer (BM) model. The first model (BC) preferentially selected the action that it believed was associated with the controlled machine (i.e., the model made choices based on a control belief, see **Figure 4**, top panel) while the second model (BM) preferentially selected the action associated with the best-rewarding Gaussian irrespective of whether this Gaussian was or was not associated with the controlled machine (i.e., the model made choices based on the magnitude of the reward; see **Figure 4**, bottom panel).

**Figure 4.**
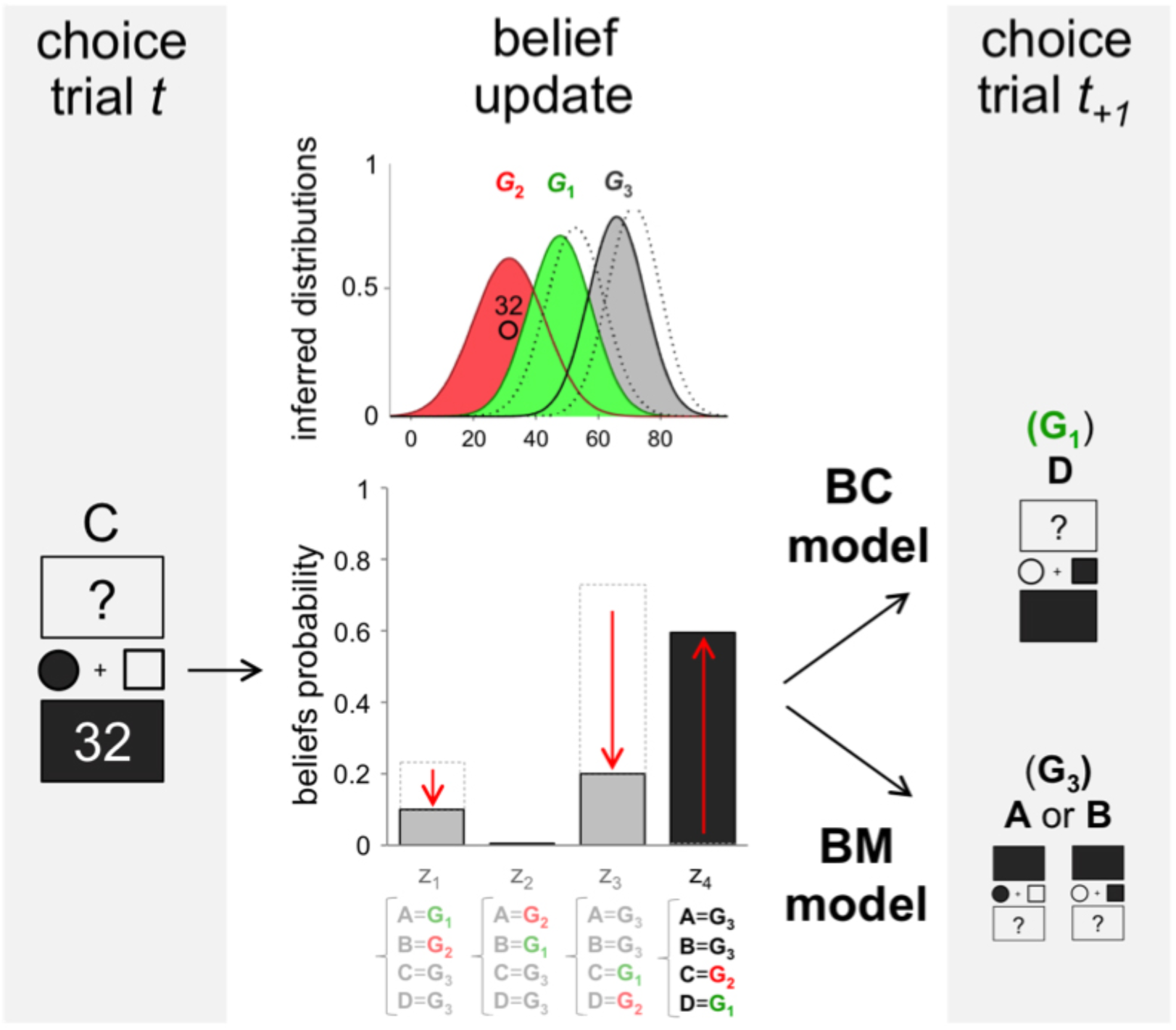
Schematic of the two-stages decision process in BC and BM models. On trial *t*, the circle button and the bottom machine are chosen, and ‘32’ is obtained as a reward. Both BC and BM models infer the current state of the world (*Z*_1_, *Z*_2_, *Z*_3_, or *Z*_4_, bottom panel) based on the inferred reward distributions (*G*_1_, *G*_2_, *G*_3_, top panel), the volatility parameter and the past history of actions and rewards. The models also update the mean and precision of the three underlying distributions (from dashed to solid distributions, top panel). Because the two models aim to maximize different statistics (control for BC, value for BM), they end up choosing different actions from the state inferred (here, *Z*_4_). Thus, on the next trial, the BC model chooses the left button and the bottom machine (**D** = the best-rewarding action of the *controlled* machine), while the BM model chooses either **A** or **B**, i.e., the current *best-rewarding* buttons.

### Action selection

Across all four models, action and belief values were used to drive action selection. For each trial, this selection was made through a *softmax* rule based on either updated action-values or beliefs (Daw *et al*., 2006). Under this rule, one action is stochastically selected according to the difference between each action’s expected value:

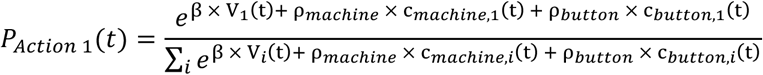

where *i* enumerates over all possible choices and *c_machine_*_,1_ and *c_machine_*_,1_ were defined as the stickiness to the previous choice irrespective of the reward history:

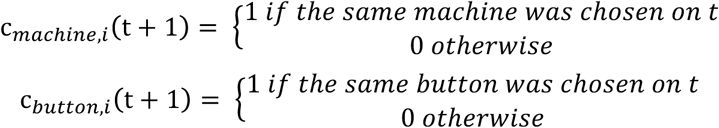

The exploitation intensity parameter β is fitted and represents the strength of the action values or beliefs on action selection. The parameters ρ*_machine_* and ρ*_button_* capture the participant’s propensity to perseverate with their action choice, which cannot be explained by reward history (Lau and Glimcher, 2005).

### Parameter fitting

Model parameters were fitted based on participants’ actions. Model fitting was performed separately for each participant and each condition. The best parameters were those maximizing the log-likelihood (LLH), which is defined as the sum of the log of the model’s fit to participants’ action choices. Thus, LLH close to 0 indicates a good model fit. To test the different possible combinations of parameters, we used a slice sampling procedure (Bishop, 2006). More specifically, using three different starting points drawn from uniform distributions for each parameter, we performed 100,000 iterations of a gradient ascent algorithm to converge on the set of parameters that best fit the data.

All four models shared the same three parameters: the perseveration biases ρ*_machine_* and ρ*_button_*, and the exploitation intensity parameter β. The two Bayesian models (BC and BM) had no additional parameter to fit since the parameters used to compute the beliefs were inferred. The RL and CF models shared the learning rate parameter α*_F_*, but the CF model had 4 additional parameters, including the three counterfactual learning rates and the reference point (*P*). To account for the risk of overfitting, a relative quality-of-fit metric, the Bayesian Information Criterion (BIC), was also computed. The BIC penalizes models with a high number of parameters:

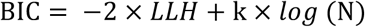

with *k* being the number of parameters and *N* the number of trials.

BIC values were compared between our four models (RL, CF, BM and BC). As an approximation of the model evidence, individual BICs were fed into the MBB-VB toolbox (Daunizeau, Adam and Rigoux, 2014), which is a procedure that estimates how likely it is that a specific model generates the data of a randomly chosen subject (the posterior probability of a model, PP) as well as the probability that a given model fits the data better than all other models in the set (exceedance probability, XP).

### Choice simulation

The four resulting models (RL, CF, BM and BC) were simulated with the best-fitting parameters, and they underwent the same experimental conditions as participants did. For each trial, the outcome given to the model was the one associated with the model’s choice, and not the participant’s choice. Simulations were used to provide aggregated measures of model performance (e.g., **Figure 5B**) but also to compare trial-by-trial choice sequence after reversal across models (e.g., **Figure 6A**).

## Experiment 1

### Method

#### Participants

Sixteen participants (8 females, with ages between 20 and 33 years old) took part in Experiment 1. They provided written informed consent prior to the experiment and were all paid 20 euros for each experimental session that was completed. No participants had a history of neurological or psychiatric disorders, and all had a normal or corrected-to-normal vision. The experiment was approved by the local ethics review board (CCP C07–28). Participants were informed about the general procedure of the experiment through detailed written instructions (see **Supplementary Information**).

#### Stimuli and trial structure

On one half of the experiment, the first choice consisted in selecting a machine, then selecting a button, and the reverse occurred for the other half (button first, then machine). The order of choice was counterbalanced within participants.

When the first choice was about the machine, a typical trial started with the presentation of two machines above and below a central fixation. Each machine was filled with a question mark (see **Figure 2A**, top panel, “machine first”). Participants had 700 ms to make their choice. Once the selection made, the question mark within the chosen machine disappeared. After a 500ms delay two buttons (a square and a circle) appeared on both sides of the central fixation. Again, the participant had 700 ms to choose one button by pressing the corresponding key. The chosen button was then filled with white to confirm the participant’s key press. Once the choice is made, the two slot machines were spun for 200 ms. The gain corresponding to the chosen button then appeared in the chosen machine for 800 ms. If the participants did not press a key within the 700 ms delay, or if the wrong key was pressed, the trial was “missed”, and the next trial started. Each trial lasted approximately 3 s.

The same timeline applied to trials in which the first choice to make was about selecting a button (**Figure 2A**, bottom panel, “button first”). As mentioned above, the spatial mapping of task stimuli and response keys was counterbalanced across the 16 participants: for half of them the machines were positioned on a vertical line, whereas the buttons were on a horizontal line (as represented in **Figure 2A**). This mapping was reversed for the other half, as were the response keys (see **Figure S1, Supplementary Information**).

Participants were informed that they would always win the sum of the gains from both machines on each trial. Thus, every 208 trials, a feedback screen displayed the participant’s current payoff, which was graphically represented as the sum of the average gains produced by each machine during the last 208 trials. In total, a session consisted of 832 trials. Each session was preceded by a short training (64 trials).

#### Experimental sessions

Participants completed 3 sessions, and each one was carried out on a different day and lasted approximately one hour. Each session required participants to track occasional changes in the structure of the task environment and to adjust their choices according to whether these changes related to either i) the *statistical dependency* between the option chosen and the subsequent outcome, or ii) the *value* or iii) the *variability* of the outcomes produced by each machine.

Thus, each session was defined according to the type of statistic manipulated in the task:

A. The “**statistical dependency**” between the action made and the resulting outcome was manipulated in the first experimental session. This session implemented a “controlled” (divergent) machine for which each button led to a different outcome, and a “non-controlled” (non-divergent) machine for which the reward was the same, regardless of the button that was pressed. For the controlled machine the gains associated with the best- and least-rewarding buttons were discretized rewards drawn from Gaussian probability distributions with identical variance (SD) but different means (see **Figure 5A**, **left panel**, green and red distributions, means = 58 and 42, SD = 10, respectively). For the non-controlled machine, the gains associated with both buttons were drawn from the same Gaussian (**Figure 5A, left panel**, grey distribution, mean = 50).

B. The “**value**” of each machine was manipulated in the second session by implementing a machine that was on average more rewarding than the other while keeping the two machines non-divergent. Thus, regardless of the button pressed, outcomes from each machine were drawn from Gaussians with identical variance but different means, such that the mean “value” of one machine (the “best-rewarding machine”, see **Figure 5A**, **middle panel**, light grey, mean = 58, SD = 10) was systematically higher than the other (the “least-rewarding machine”, dark grey, mean = 42, SD = 10).

C. In a third session the “**variance**” of each machine was manipulated by making the gains from one machine more variable than the other while keeping the two machines non-divergent. Thus, regardless of the button pressed, the two machines were associated with Gaussian distributions that had the same mean but different variance (low- and high-variable machines: mean = 50, SD = 5 and 15, respectively) (see **Figure 5A, right panel**).

We were first interested in assessing (i) whether, and how, the three statistics manipulated could influence participants’ control beliefs, and second (ii) whether and how well each class of models could account for this influence on participants’ choice behaviours. To independently assess the influence of the “value” and “variance” statistics on choice, the last two sessions did not implement any “divergent” machines. As a result, participants had no real control over the gains produced by the machines. The reason for this was twofold. First, it allowed for assessing whether choice behaviours modified in situations where one was told that events in the task were under one’s own control but where no true control in fact existed – such as in classical settings implementing the so-called “illusion of control” (Stefan and David, 2013). Second, the procedure allowed for testing how best fitting models –i.e., models that best accounted for participants’ choice under normal conditions– performed in a situation of *illusory control*, and how well these models effectively accounted for the participant’s data in this situation.

Finally, to keep all sessions as similar as possible, the same instructions were delivered across all three sessions. Thus, instructions in the “value” and the “variance” sessions were the same as those given in the “dependency” session, which meant that participants were not told they had no control over the machines in these conditions. All participants always started with the “dependency” session that implemented divergent and non-divergent machines followed by the value and variance sessions in counterbalanced order across participants.

#### Reversals

Each session comprised 32 “episodes”. An episode referred to an uninterrupted series of trials before a reversal occurred. The number of trials within an episode was on average 26 but varied between 14 and 38 (uniformly jittered) to make reversals as unpredictable as possible. In the “action-outcome dependency” session, two types of reversal could occur: either the buttons or the machines reversed such that the controlled machine became the non-controlled machine, or the best-rewarding button became the least-rewarding button. For the value and variance sessions, only non-divergent machines were implemented, so that only “machine” reversals occurred: either the best-rewarding machine became the least-rewarding machine (“value” session) or the low-variable machine became the high-variable machine (“variance” session).

#### Modelling

To simulate participants’ choices, we implemented the same four models that were previously described (RL, CF, BC and BM). To test if participants would adapt their strategy to the session, we fitted the models’ parameters separately across the three different experimental sessions. As mentioned above, in both the “value” and “variance” sessions, instructions were the same as those delivered in the “dependency” session: participants were not told they had no real control over the machines. This was explicitly accounted for by in the two generative models (BC and BM) through implementing the same latent states (i.e., generative distributions) as in the “dependency” session. Thus, our two generative models assumed there were a controlled and a non-controlled machine for all conditions.

**Figure 5.**
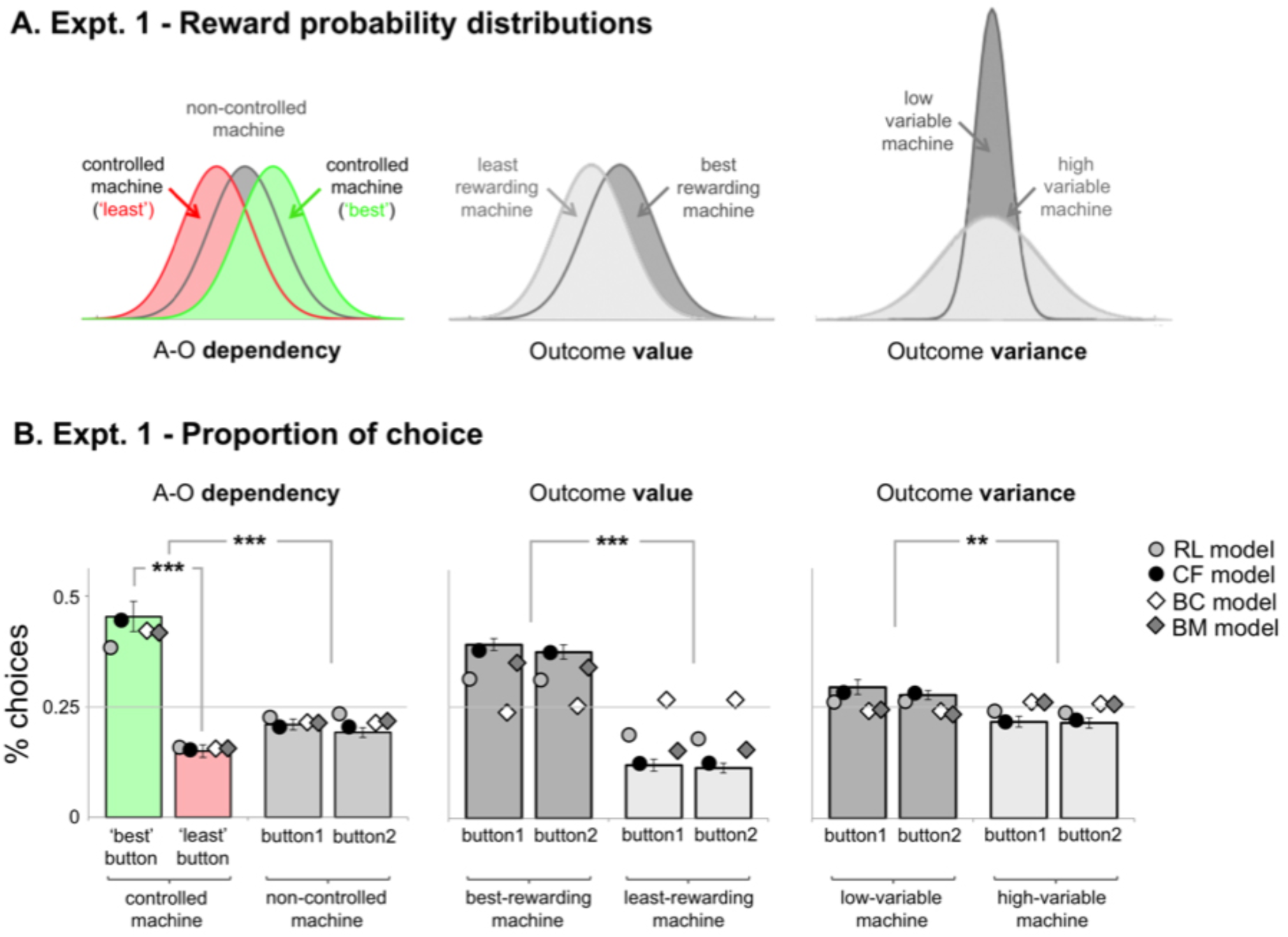
(A) Reward probability distributions associated with each button and machine of each experimental session from Experiment 1. *Left panel*: In the first session, action-outcome dependency was implemented for one machine only. For this “controlled” machine, the outcome depended on the button choice: one button led to a mean outcome of 58 (green) whereas the other button led to a mean outcome of 42 (red). For the other “uncontrolled” machine (dark grey), the outcome displayed was drawn for a *unique* Gaussian distribution, irrespective of the button being chosen. *Middle panel:* In the “value” condition, no machine was controlled, but one bandit was best rewarded (dark grey, mean outcome: 58) than the other (light grey, mean outcome: 42). *Right panel:* In the “variance” condition, the mean outcome (50) was the same for both machines, irrespective of the button chosen, but outcomes from one machine were more variable than outcomes from the other machine (light grey, SD = 15, *vs*. dark grey, SD = 5). **(B) Mean proportion of choice for the three sessions, and for each button and machine**. Bars: participants’ choices (%); dots and diamonds: models’ choices (%). RL: reinforcement-learning model; CF: counterfactual model; BC: Bayesian-controller model; BM: Bayesian-maximizer model. The horizontal grey line indicates chance level (0.25%). All error bars indicate standard error. For the sake of visibility, models’ error bars are not shown. Three-stars: *p* < 0.001.

### Results

#### Percentage of choices

We first assessed whether subjects could discriminate between the two (divergent) states of the *controlled* machine relative to the non-controlled machine by comparing the choice proportion for each button of each machine within each session. As expected, participants discriminated well between the two buttons of the controlled machine in the dependency session (best- and least-rewarding buttons: 0.45 vs. 0.15, t(15)= 6.3, *p* < 0.001, **Figure 5B**, green vs. red bars, left panel), while choosing button 1 and button 2 of the non-controlled machines equally in all sessions (all t’s < 1.62, all *p*’s > 0.12; **Figure 5B**, grey bars). We then compared button preferences *across* all sessions. To do so, we subtracted choice proportion for one button from choice proportion for the other button within each preferred machine and compared the difference across sessions using a one-way ANOVA (dependency vs. value vs. variance). The ANOVA confirmed that “button” preferences differed across the 3 sessions (F(2,45)=28.98, *p* < 0.001, 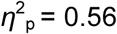). Thus, participants discriminated between buttons of the preferred machine in the *dependency* session to a far greater extent than in the *value* and *variance* sessions (0.30 vs. 0.016, and 0.30 vs. 0.017, respectively, post hoc tests: all *p*’s < 0.001).

Second, we tested whether participants showed a preference for one machine over another within each session by comparing choice proportion for each machine against the chance level (0.50). We found that participants showed a marked preference for the *controlled* machine in the dependency session (0.62 vs. 0.50, t(15) = 4.86, *p* < 0.001) as well as a marked preference for the *best-rewarding* (0.76 vs. 0.50, t(15) = 10.67, *p* < 0.001) and the *low-variable* (0.57 vs. 0.50, t(15) = 3.04, *p* = 0.004) machines in the value and variance conditions, respectively. Finally, we compared the proportion of choice for the preferred machine across all 3 sessions. The one-way ANOVA revealed that “machine” preferences differed across the 3 sessions (F(2,45)=21.31, *p* < 0.001, 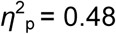), Thus, participants chose the *best-rewarding* machine (“value” session, 0.76) more than the *controlled* machine (“dependency” session, 0.62), and both the *controlled* and *rewarding* machines were chosen more than the *low-variable* machine (“variance” session, 0.56) (post hoc tests, all *p’s* < 0.05).

Note that, in all sessions, participants were able to quickly adjust to machine and/or button reversals: on average, the plateau of performance was reached within 5–10 trials after reversal (see **Figure 6A**, “reversal learning curves”).

#### Model comparison

Participants’ trial-by-trial choice sequence were best accounted for by the CF model than by all other models in the set (RL, BM or BC). This was true for all conditions (exceedance probability > 98%) (**Table 1 and Figure 6B**). In addition to comparing model parameters across conditions and subjects, we also evaluated the generative performance of each concurrent model, i.e., its ability to replicate the participant’s proportion of choices as well as the participant’s trial-by-trial choice sequence after reversal (Palminteri *et al*., 2017). To do so, the 4 models were simulated with the best-fitting parameters for the whole experiment. Crucially, only the CF model showed a pattern of choices similar to participants in *all* sessions either with regard to the choice of the machine or to the choice of the button (see **Figure 5B**, CF = black circle).

Then we plotted the model learning dynamics before and after a reversal. Again, only the CF model was as flexible as participants and adjusted to reversals with a similar dynamic (see **Figure 6A**, CF = red bars). In the “dependency” session, more specifically, the CF model outperformed all 3 competitors for both types of reversals. Thus, CF was able to retrieve the “controlled” machine and the “best-rewarding” option as quickly as participants, while the 3 other models adjusted more slowly, as particularly evidenced by the RL model after a button reversal (**Figure 6A**, top panel). In the “value” session, CF also better simulated the participants’ choices than all competing models (**Figure 6A**, bottom panel, left). Note that the BC model (“Bayesian-controller”, dark green bars) aimed to maximize control, i.e., preferentially chose the option associated with the controlled machine. Thus, its poor performance in this session with no true control was no surprise. The same trend applied to the “variance” session where no machine was controlled either (**Figure 6A**, bottom, right). In this session, human participants showed a marked preference for the “low-variable” machine and switched their choice after reversal to retrieve this preferred machine. Importantly, only the CF model was able to simulate this preference for poor variable choice outcomes.

**Figure 6.**
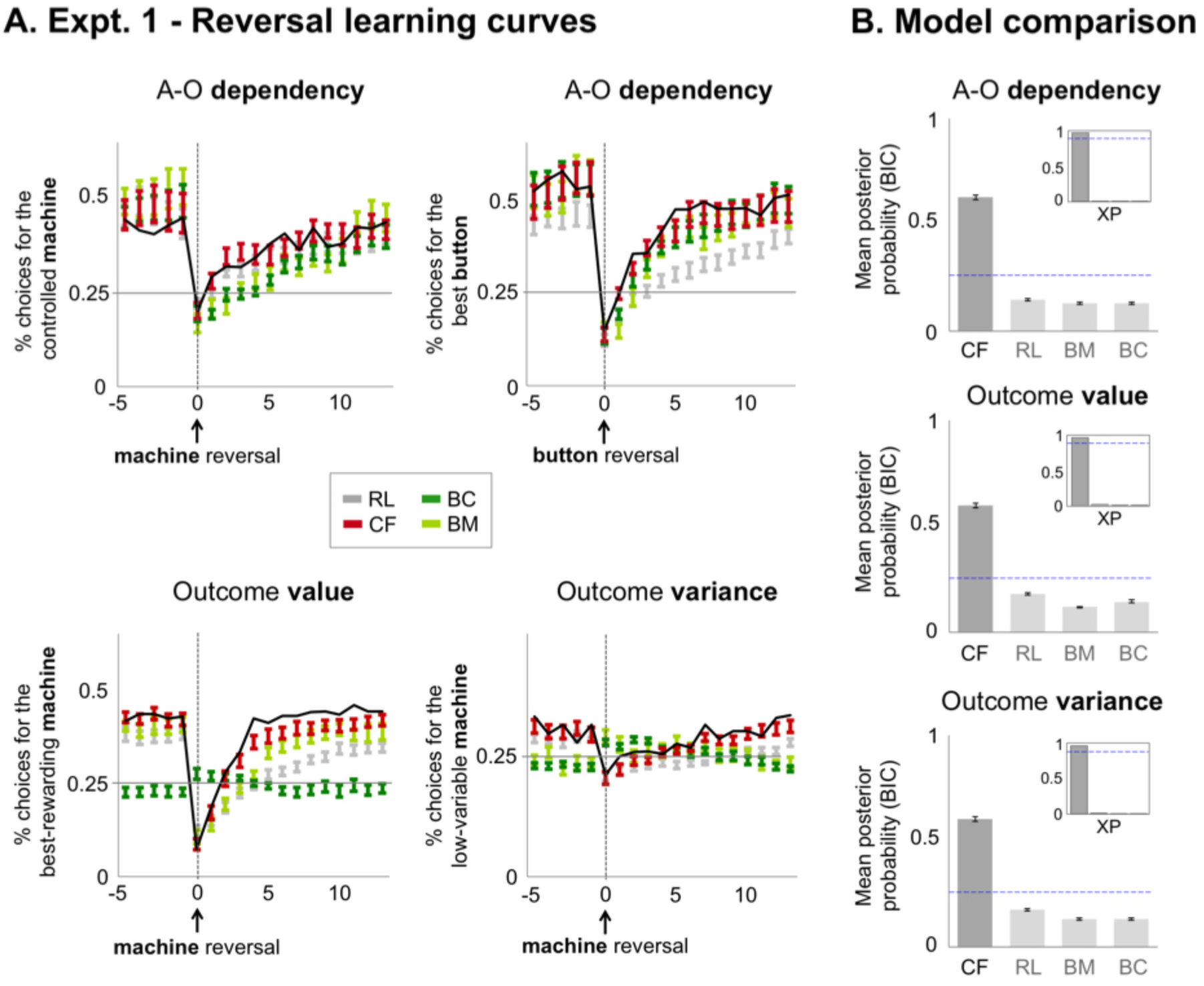
(A) Reversal curves for human participants (solid black line) and models (colored bars) up to 15 trials after a “machine” or a “button” reversal. *Top panel:* reversal curves for the A-O dependency session, after a (controlled vs. non-controlled) machine reversal, or a (best vs. least-rewarding) button reversal. *Bottom panel:* reversal curves for the value and the variance session after a machine reversal (best vs. least-rewarding machine, or low vs. high variable machine, respectively). For the sake of readability, subjects’ error bars are not shown. Model simulations: CF (red bars); RL (light grey); BM (light green bars); BC (dark green bars). Bars indicate standard error. RL: reinforcement-learning model; CF: counterfactual model; BC: Bayesian-controller model; BM: Bayesian-maximizer model. Dashed vertical lines indicate reversal point. Horizontal grey lines indicate chance level. **(B) Comparison of the posterior probability (PP) of each model, for each session**. The PP is calculated from the BIC, which penalizes model complexity. The blue dashed line represents the chance level at 0.25. The insert chart shows the exceedance probability (XP) of each model in the set. The blue vertical dashed line shows the 95% threshold. In all three sessions, the CF model best explained the data.

### Experiment 1: Preliminary discussion

The first experiment tested whether, and how well, human participants adjusted to self- vs. externally generated changes in a task in which the source of these changes was uncertain.

Our results show that participants discriminated well between best- and least-rewarding buttons as well as between controlled and non-controlled machines. Hence, they preferentially chose the controlled machine over the non-controlled machine while exhibiting a marked preference for both highly rewarding and low-variable machines. In the context of goal-directed control, this preference for high reward and low variance is reminiscent of the literature on self-attribution biases: adults are more likely to believe they control the occurrence of *positive*, relative to negative, events (e.g., Mezulis *et al*., 2004) while spontaneously assuming that series of *low-variable* events are more likely to be generated by intentional than non-intentional agents (e.g., Boland and Pawitan, 1999; Caruso, Waytz and Epley, 2010). Unsurprisingly, the pattern of preference exhibited across all 3 sessions suggests that participants construe their action, not only as a mean to make a difference in the world (instrumental divergence) but also as an instrument to bring about positive events and to reduce the inherent variability of the environment.

Both quantitative (BIC) and qualitative (simulated learning curves) results showed that a model drawing on pure associative processes (RL) cannot fully explain participants’ behaviours and neither can a generative model that makes choices based on either gain (BM) or control (BC) maximization strategies. Rather, we found that a model (CF) deriving the consequences of the forgone action from the current action taken and assuming relative (i.e., context-dependent) divergence between both best explained the data.

While BC and BM models had explicit priors for control in the task – assuming distinct outcome distributions depending on the subject’s choice –, the CF model was endowed with a more general prior about instrumental divergence. This prior implements the belief that taking a specific action (e.g., choosing option A vs. B) makes a difference in terms of the outcome. Importantly, instrumental divergence is a reliable proxy for goal-directed control as the greater the action *diverges* with respect to its contingent states (the factual and counterfactual outcomes), the more flexible control one has over the environment. The fact that CF best explains data in all conditions suggests that human subjects construe their causal power based on such a prior. Interestingly, the CF model also best accounted for the participants’ choice even when no true control existed, which suggests that this prior holds as a default belief, whereby goal-directed actions are thought to be causally efficient (i.e., divergent) in nature.

This study had two limitations. First, all sessions were not fully counterbalanced between subjects. All participants underwent the “dependency” session first, and then the two remaining sessions in which no true control was implemented. Additionally, instructions given across all three sessions systematically emphasized the notion of control over the task. In a follow-up experiment (20 subjects, 3 one-hour sessions each), we thus ran a similar task while carefully controlling for these two potential biases. Sessions were fully counterbalanced and both verbal and written instructions were kept as minimal as possible (see **Appendix B**, “Experiment 1b: Methods and Results”).

Most of the results from experiment 1 were replicated. As expected, participants exhibited a strong preference for high rewards and preferentially chose low-variable machines, regardless of their overall value. We also found that participants were able to discriminate between causally efficient actions and to identify *where* in the task environment choosing one action rather than another made a difference to the outcome (the controlled machine), and where it did not (the non-controlled machine). Finally, we again found that a model based on a simple context-dependent counterfactual rule (CF) fitted the data better than all competing models, including a pure reinforcement learner (RL) and a model that explicitly aimed at maximizing reward through Bayesian inference (BM) (**Appendix B**).

In both experiment 1 and its follow-up, each statistic (dependency, value, variance) was tested within a different session, therefore limiting the opportunity to test and control for their potential interactions. In a second experiment, we addressed this limitation by implementing a factorial design in which these statics were systematically crossed. In addition to controlling for interaction effects, this experimental design allowed for better characterization of subjects’ choices in situations where these statistics were explicitly conflicting.

**Table 1.** Mean posterior probability (±SEM) of each model (*PP*) in each session and/or experiment. The exceedance probability (XP) refers to the probability that a given model fits the data better than all other models. CF: counterfactual model; RL: reinforcement-learning model; BC: Bayesian-controller model; BM: Bayesian-maximizer model.

| Session |  | CF |  | RL |  | BM |  | BC |  |
| --- | --- | --- | --- | --- | --- | --- | --- | --- | --- |
|  |  | *PP* | *XP* | *PP* | *XP* | *PP* | *XP* | *PP* | *XP* |
| Expt. 1a | Dependency | 0.60<br>($\pm 0.011$) | 99% | 0.14<br>($\pm 0.005$) | < 1% | 0.12 ($\pm 0.003$) | < 1% | 0.12 ($\pm 0.003$) | < 1% |
| | Value | 0.57<br>($\pm 0.011$) | 98% | 0.17<br>($\pm 0.013$) | < 2% | 0.11<br>($\pm 0.003$) | < 1% | 0.14<br>($\pm 0.006$) | < 1% |
| | Variance | 0.58<br>($\pm 0.011$) | 98% | 0.16<br>($\pm 0.006$) | < 2% | 0.12<br>($\pm 0.005$) | < 1% | 0.12<br>($\pm 0.005$) | < 1% |
| Expt. 1b | Dependency | 0.47<br>($\pm 0.009$) | 92% | 0.19<br>($\pm 0.006$) | < 4% | 0.17<br>($\pm 0.005$) | < 3% | 0.17<br>($\pm 0.005$) | < 3% |
| | Value | 0.53<br>($\pm 0.009$) | 97% | 0.18<br>($\pm 0.006$) | < 2% | 0.14<br>($\pm 0.005$) | < 1% | 0.13<br>($\pm 0.004$) | < 1% |
| | Variance | 0.50<br>($\pm 0.009$) | 96% | 0.17<br>($\pm 0.005$) | < 2% | 0.16<br>($\pm 0.005$) | < 2% | 0.15<br>($\pm 0.005$) | < 2% |
| Expt. 2 | | 0.85<br>($\pm 0.004$) | 99% | 0.06<br>($\pm 0.002$) | < 1% | 0.04<br>($\pm 0.001$) | < 1% | 0.04<br>($\pm 0.001$) | < 1% |

## Experiment 2

### Method

#### Participants

Twenty-six participants (14 females, age between 21 and 40 years old) took part in Experiment 2. As before, they provided written informed consent prior to the experiment and were all paid 80 euros for the whole experiment (4 sessions). No participants had a history of neurological or psychiatric disorders, and all had a normal or corrected-to-normal vision. The experiment was approved by the local ethics review board (CCP C07–28). Participants were informed about the general procedure of the experiment through detailed written instructions.

#### Experimental sessions

The task (stimuli, timeline, and trial structure) was identical to the one used in experiment 1. The only difference was implemented by the experimental design. Each participant completed 4 sessions, and each lasted approximately one hour. Each session was carried out on a different day. A session consisted of 9 experimental conditions with 140 trials each (**Figure 7**) and was preceded by a brief training (64 trials). The order of conditions was pseudo-randomized within each session.

As in the previous experiment, the “statistical dependency” between action and outcome was manipulated by implementing a controlled (divergent) machine for which each button led to a different outcome, and a non-controlled (non-divergent) machine for which the reward was the same, regardless of the button that was pressed. For the controlled and non-controlled machines, the gains associated with each button were discretized rewards drawn from Gaussian probability distributions, whose variance and mean depended on the condition (see below). The “value” and “variance” dimensions were crossed within a 3-by-3 factorial design, with each dimension varying across three levels (**Figure 7**):

1. The “**value**” dimension referred to the mean of the reward probability distribution associated with each machine. The mean of the controlled machine could vary across three different values (i.e., low = 42, medium = 54, high = 62), whereas the mean of the non-controlled machine was kept constant (i.e., 62). Low, medium, and high values characterized the average reward delivered by the controlled, *relative* to the non-controlled, machine (**Figure 7**, x-axis). The “low” value level indicated that the controlled machine was less rewarding on average than the non-controlled machine, whereas the “high” level indicated that the non-controlled machine was more rewarding than the non-controlled one.

2. The “**variance**” dimension referred to the standard deviation (SD) of the reward probability distribution associated with each machine. The standard deviation of the controlled machine was kept constant throughout the task (SD = 10) whereas the standard deviation of the non-controlled machine varied across three levels (low, SD = 5; medium, SD =10; high, SD = 15) (**Figure 7**, y-axis). These 3 levels characterized the variability of the rewards delivered by the controlled, *relative* to the non-controlled, machine. Thus, the “low” variance level indicated that the controlled machine was less variable than the non-controlled machine, whereas the “high” variance level indicated that the controlled machine was more variable than the non-controlled one.

Because the 3 levels of each dimension characterized the value and variance of the controlled machine *relative* to the non-controlled machine, we now refer to these as low, medium, and high “relative levels”. In the following, we looked at whether choice proportion changed as a function of the relative value and relative variance of the controlled machine. Specifically, we asked whether the proportion of choice for the best-rewarding button and/or the controlled machine would change as the controlled machine became more or less rewarding, or more or less variable, than the non-controlled machine.

#### Reversals

Finally, button or machine reversals could occur within each experimental condition as described before. Button reversals consisted of the best-rewarding button (e.g., the square) becoming the least-rewarding button (e.g., the circle) for the controlled machine, whereas machine reversals consisted in the controlled machine becoming the non-controlled machine. Within each experimental condition, 6 reversals (3 machine reversals and 3 button reversals) could occur after a variable number of trials (between 14 and 26, uniformly jittered).

**Figure 7.**
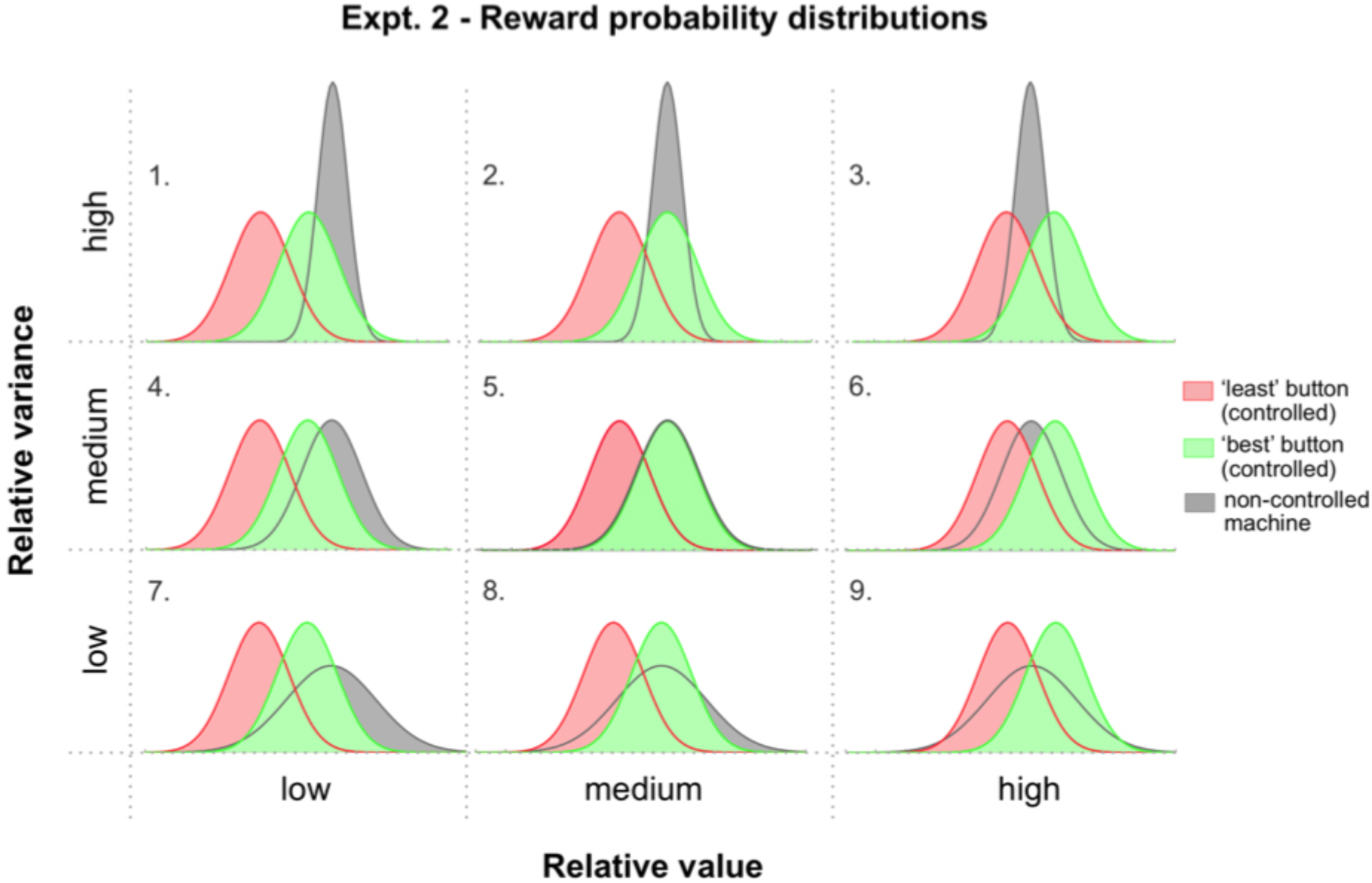
Schematic of experimental conditions in Experiment 2. Contribution of action-outcome dependency (controlled vs. non-controlled machine), outcome value (x-axis) and outcome variance (y-axis), to the participant’s choice, was assessed by manipulating the reward probability distributions associated with each button of each machine. Red and green Gaussian distributions: rewards from the controlled machine when the best- and least-rewarding buttons were selected, respectively. Grey distributions: rewards from the non-controlled machine, irrespective of the button selected. *X-axis*: the value of the controlled, *relative* to the non-controlled, machine, varied across three levels (low, medium, and high) – e.g., “low” level. the value of the controlled machine was low relative to the non-controlled machine. *Y-axis*: the variance of the controlled, *relative* to the non-controlled, machine, varied across three levels (low, medium, and high) – e.g., “low” level: the variance of the controlled machine was low relative to the non-controlled machine. Experimental conditions are numbered from 1 to 9 (top left to bottom right).

### Results

#### Percentage of choices

We investigated the effect of the value and the variance dimensions, together with their interaction, on two dependent variables: (i) the proportion of choice for the *controlled machine*, and (ii) the proportion of choices for the *best-rewarding button* of the controlled machine. As in the previous experiment, the proportion of choice for the best-rewarding button was normalized by subtracting the proportion of choice for the least-rewarding button from it within each condition.

The proportion of choices for the controlled machine as well as the proportion of choices for the best-rewarding button were analysed using two 3 × 3 repeated-measures ANOVAs with the value (low vs. medium vs. high) and variance (low vs. medium vs. high) as within-subjects factors. Participants discriminated well between the controlled and non-controlled machines across all 9 conditions, but the proportion of choice for the controlled machine differed significantly as a function of the dimension manipulated. Thus, we found a significant main effect of the value (F(2,50)=283.50, *p* < 0.001, 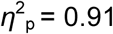) and a significant main effect of the variance (F(2,50)=5.48, *p* = 0.007, 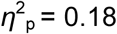) factor on the proportion of choice for the controlled machine. The proportion of choice for the controlled machine progressively increased as its relative value increased (low < medium < high, post hoc test, all *p*’s < 0.001), but also increased when its relative variance *decreased* (low vs. medium, *p* = 0.009; medium vs. high, *p* = 0.04). These results are consistent with the high-value and low-variance biases observed in experiment 1, in which participants tended to preferentially select the machine with the highest value and the lowest variance (see **Figure 5**).

The value-by-variance interaction effect was also significant (F(4,100)=6.66, *p* < 0.001, 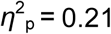). Thus, when the relative value of the controlled machine was high, participants more often chose this machine irrespective of the variance dimension; that is, they chose the controlled machine in similar proportions whether the controlled machine was highly or poorly variable (post hoc tests comparing low vs. medium vs. high variance, all *p*’s > 0.12). On the other hand, when the value of the controlled machine was low, participants tended to choose the controlled machine more when it was poorly, rather than highly, variable (comparing low vs. medium variance, *p* = 0.07; low vs. high variance, *p* = 0.005). In other words, the variance dimension had the strongest effect on the choice of the controlled machine when the *value* of this machine was the *lowest* (**Figure 8**, **top panel**, “Machine choice”, “LOW value”).

Similar to experiment 1, we then compared the proportion of choice for the best-rewarding button *across* conditions. We again found significant main effects of the value (F(2,50)=78.81, *p* < 0.001, 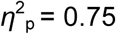) and variance (F(2,50)=4.28, *p* < 0.019, 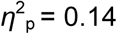) dimensions. With respect to the value dimension, the higher the relative value of the controlled machine, the more often participants chose the best, relative to the least, rewarding button (post hoc tests: low vs. medium = 0.28 vs. 0.17, *p* < *0.001*; medium vs. high, *p* < 0.001). In other terms, the more rewarding the controlled machine was, relative to the non-controlled machine, the more participants discriminated between each button, and the more their choice reflected the true “divergence” of the controlled machine (**Appendix C, Figure S2**). The same was observed for the variance dimension: the proportion of choice for the best, relative to the least, rewarding button increased as the relative variance of the controlled machine *decreased* (post hoc tests comparing low vs. medium variance: 0.15 vs. 0.17, *p* = 0.07; medium vs. high variance: 0.17 vs. 0.19, *p* = 0.005) (**Appendix C, Figure S3**). Finally, the value-by-variance interaction effect was also significant (F(4,100)=8.54, *p* < 0.001, 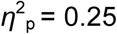). We found the same pattern of interaction for the machine choice: the variance dimension had the strongest effect on button choice as the value of the controlled machine *decreased* (**Figure 8, bottom panel**, “Button choice”, “LOW value”).

In sum, for both dependent variables (machine and button choices), the outcome value had an overwhelming influence on participants’ choice and this influence largely overrode the effect of variance. As a consequence, the effect of variance could only be observed in conditions in which the value of the controlled machine was the lowest (see **Figure 8, top** and **bottom panels**)

**Figure 8.**
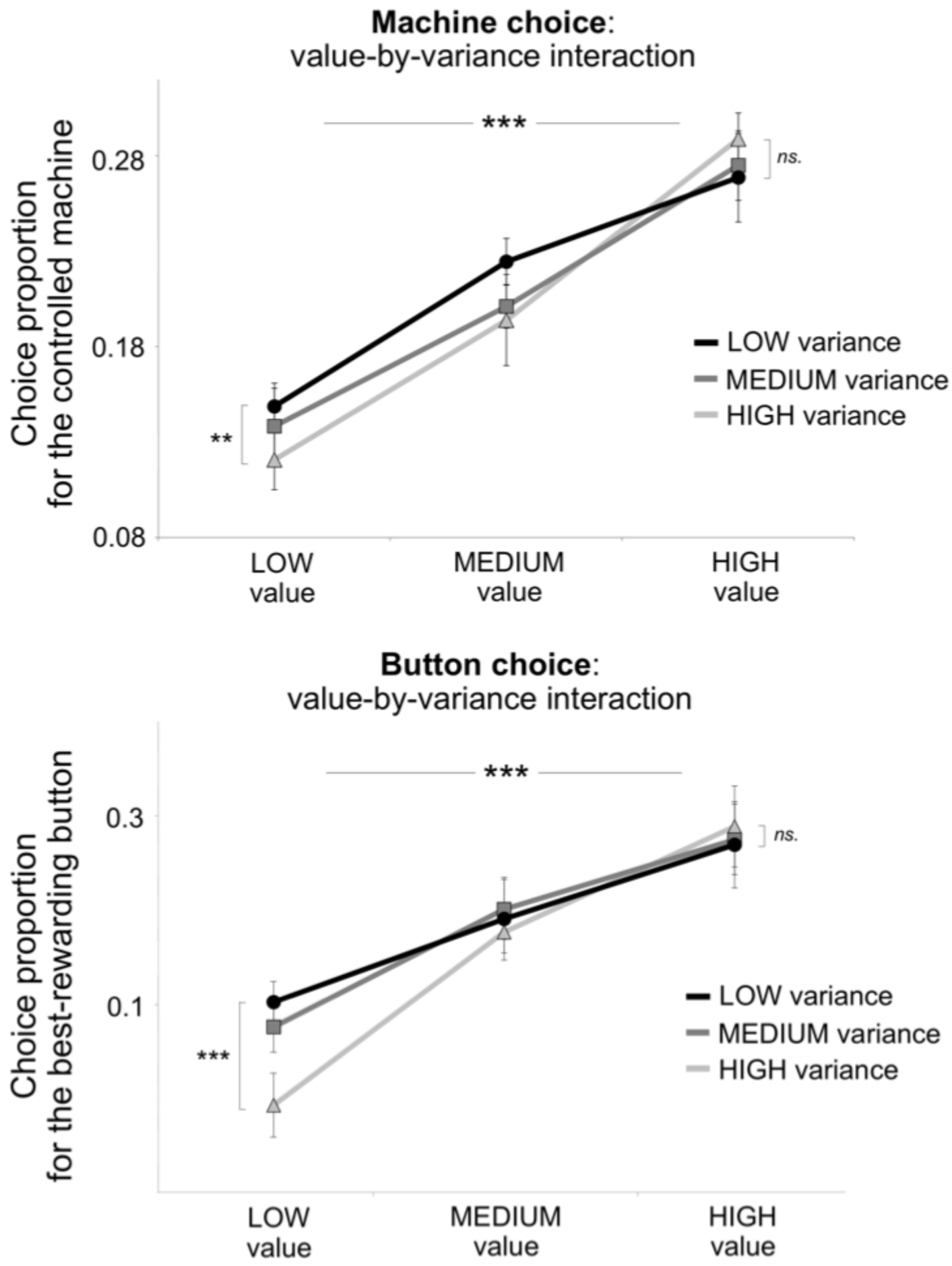
Participants’ performance: mean proportion of choice (±SEM) across each dimension manipulated. *Top panel:* proportion of choice for the controlled machine in trials in which the machine had a low, medium, or high *value*, relative to the non-controlled machine (*x*-axis), and had a low, medium, or high, *variance*, relative to the non-controlled machine (black, dark grey, and light grey, solid lines, respectively). The interaction effect between the value and variance factors was significant: the *variance* dimension had the strongest effect on the choice of the controlled machine when the *value* of this machine was the lowest. *Bottom panel:* normalized proportion of choice for the best-rewarding button in trials in which the controlled machine had a low, medium, or high *value*, and had a low, medium, or high, *variance*, relative to the non-controlled machine. As for the choice of the controlled machine, the interaction effect between the value and variance factors was significant. Two-stars: *p* < 0.005; Three-stars: *p* < 0.001; *ns*. = *p* > 0.05.

#### Model comparison

The same four models were fitted and simulated as described before. Again, the CF model best predicted participants’ choices (exceedance probability = 99%, **Table 1**, and **Figure 9C**) with regard to choice proportion for the best-rewarding button (**Figure 9A**, CF = black circles) or to choice proportion along the value (**Appendix C**, **Figure S2**) or the variance (**Appendix C, Figure S3**) dimensions.

Importantly, the participant’s sensitivity to action-outcome dependency was best accounted for by the CF model. Thus, CF was the only model that did not underestimate the difference in choice proportion between the buttons of the controlled and non-controlled machines (see **Figure 9A**, black circle). The CF model also correctly simulated the participant’s choices along the value dimension. Thus, the CF model was able to reproduce the participant’s propensity to better discriminate between the “best” and “worst” buttons as the value of the controlled, relative to the non-controlled, machine increased (**Figure S2**). The RL (grey circles) and BM model (grey diamonds) showed a similar, although less clear-cut, pattern of choice. In contrast, the BC model (white diamonds) exhibited the same pattern of choices across all 3 levels of the dimension, and both the BC and BM models underestimated the difference between the two buttons of the controlled machine in the “high” value condition. The same applied to the “variance” dimension: both the CF and RL models were able to discriminate buttons of the controlled machine while choosing the two buttons of the non-controlled one equally often (**Figure S3**). In contrast to the BC and BM models, CF and RL also tended to choose the low-variable, relative to the high-variable, machine more often, as participants did.

**Figure 9.**
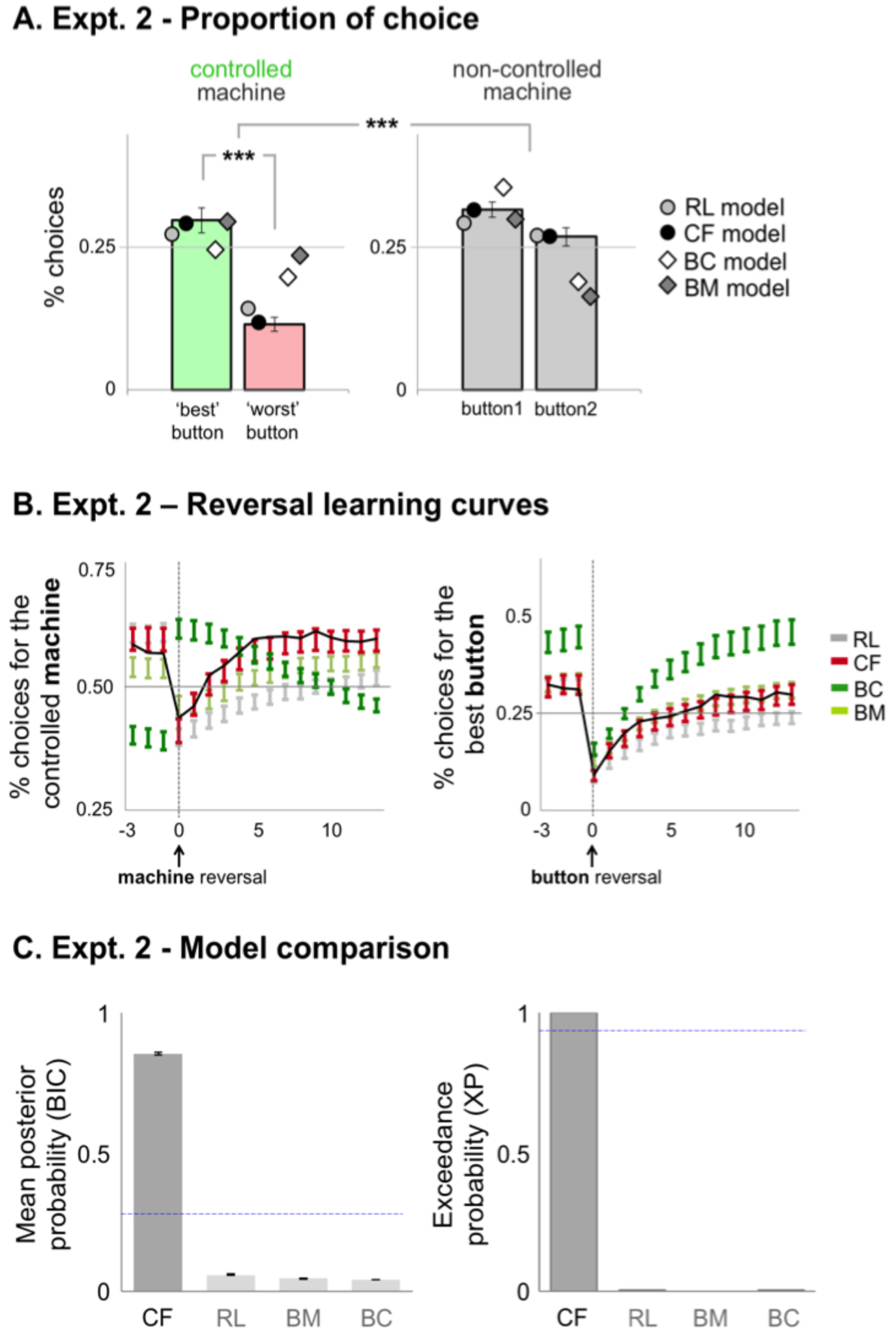
(A) Bar graphs comparing the proportion of choice (± error bars) across buttons and machines, averaged across all dimensions of the task design. Bars: participants’ performance; dots and diamonds: models’ performance. For the sake of visibility, models’ error bars are not shown. Three-stars: *p* < 0.001. **(B) Reversal curves (± error bars) for participants (solid black line) and models (colored bars)**, up to 15 trials after a button reversal. The horizontal grey line indicates chance level. Dashed vertical lines indicate reversal point (*left graph:* machine reversal; *right graph:* button reversal). Model simulation: CF (red bars); RL (light grey); BM (light green bars); BC (dark green bars). **(C) Comparison of the posterior probability (PP) of each model, for each session**. The PP is calculated from the BIC, which penalizes model complexity. The blue dashed line represents the chance level at 0.25. The right graph shows the exceedance probability (XP) of each model in the set, with the blue dashed line representing the 95% threshold.

As for participants, the models’ choices for the controlled machine and for the best-rewarding button were analysed further using two 3 × 3 ANOVAs with value (low vs. medium vs. high) and variance (low vs. medium vs. high) as within-subject factors. Relative to participants’ performance, only the CF model was able to replicate the main effects of the value (*machine choice:* F(2,50) = 289.36, p < 0.001, 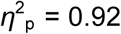; *button choice:* F(2,50) = 86.00, p < 0.001, 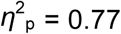) and of the variance (*machine choice:* F(2,50) = 2.50, p < 0.001, 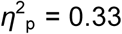; *button choice:* F(2,50) = 5.03, p = 0.01, 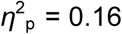) factors as well as the significant interaction effects between them (*machine choice:* F(4,100) = 21.98, p < 0.001, 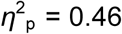; *button choice:* F(4,100) = 2.68, p = 0.035, 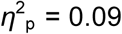; post hoc comparing low vs. high variance, *p* = 0.032) (**Figure 10**, “**CF model**”, *machine choice:* left panel; *button choice:* right panel). Importantly, none of the 3 other models could replicate this exact performance pattern.

The CF model also showed the most consistent reversal curves across conditions and outperformed all competing models when adjusting to changes according to either action-outcome dependency (**Figure 9B**), outcome value (**Appendix C, Figure S2**) or outcome variance (**Appendix C, Figure S3**). Specifically, both the BM and CF models correctly simulated the participants’ learning curves, whether in terms of dynamic (slope) and absolute performance (plateau), whereas the RL model converged to the plateau of performance more slowly than real subjects did. Note that the BC model (dark green line) was designed to preferentially choose the action associated with the controlled machine, and therefore systematically reversed choice after reversal of the best button.

**Figure 10.**
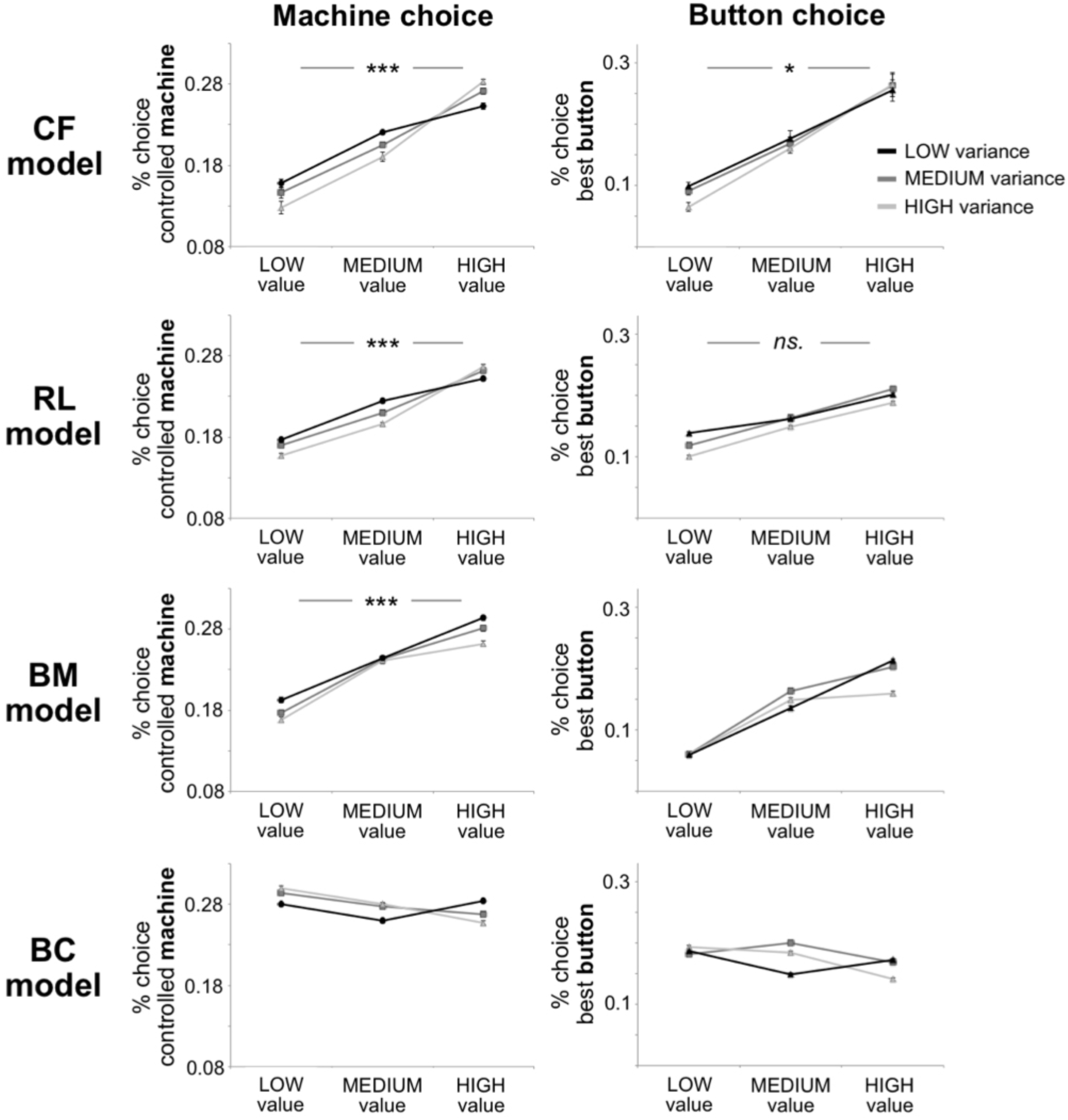
Models’ performance: mean proportion of choice (±SEM) across each dimension manipulated. Stars indicate a significant interaction between the value and variance factors. Only the CF model replicate the interactions observed in human subjects, for both machine (left) and button (right) choices. The BC model shows the *opposite* interaction effects (see Results). One-star: *p* < 0.05; Three-stars: *p* < 0.001; *ns*. = *p* > 0.05.

### Experiment 2: Preliminary discussion

Experiment 2 reproduced most of the effects that was previously obtained in a design controlling for potential interaction effects between conditions. Importantly, participants performed the task well despite no explicit available cue to signal the transition from one condition to the other. In a situation in which uncertainty was high, participants were able to monitor the different statistics implemented by the task and to adjust when these statistics changed and reversals in (either machine or button) contingencies occurred. In line with experiment 1, we found that participants chose the *controlled* machine more often when the relative *value* of this machine increased, but they also chose it when its relative *variance* decreased, in line with the literature on self-attribution biases. Likewise, when the value of control increased, participants discriminated better between the best and worst option of the controlled machine, and choice behaviour was then found to better reflect the true divergence of the controlled machine. Finally, we found a significant interaction between value and variance factors. Specifically, a significant effect of *variance* on machine and button choice was observed in *low-value* trials only. This interaction suggests that competition between both statistics is fundamentally asymmetrical. If a conflict arose, subjective preferences for highly valued options overrode preferences for options giving rise to poorly variable outcomes. On the other hand, when the difference in value between competing options was low, subjects made a choice based on variance estimates from past choice outcomes.

As expected, the overall effect of *value* on choice was well captured by algorithms that aimed to maximize reward value (RL, CF), while an optimal learner aiming to maximize control (BC) failed to account for this effect. We again found that the CF model outperformed all competitors according to both quantitative (BIC) and qualitative (reversal curves) criteria. Interestingly, CF was the only model to not systematically underestimate the difference in choice proportion between the two buttons of the controlled and non-controlled machines as well as to better discriminate between each button of the controlled machine as the value of this machine increased, as participants did. The CF model was also the only model to replicate the exact pattern of performance found in human subjects. Thus, the CF model increasingly chose the controlled machine and the best-rewarding button as the value of this machine *increased* (main effect of value) but also as its variance *decreased* (main effect of variance). Critically, choices of the CF model also exhibited a significant value-by-variance interaction effect. Thus, the effect of variance on the model’s choices was only observed in low-value trials, as in participants. Overall, we found that one single model (CF) was able to simulate participants’ performance across all three dimensions and was able to do so with the same set of parameters and same parameter values.

In the next section of the paper, we analysed and further compared the parameters of the “winning” CF model across both experiments 1 and 2, namely: (1) the reference point, and (2) the factual and counterfactual learning rates. Importantly, these two sets of parameters can be seen as direct or indirect proxies for instrumental control:

(1) The “reference” is a fitted parameter whose value approximated the mean of the reward distribution associated with the chosen action (see **Figure 11A** and **11B**). It is an add-on to the classical RL algorithm that implements control as *difference-making*. Thus, the more the value of the reference departs from the true reference, the more *divergent* actions are, and the greater the *difference* between the outcomes associated with the chosen and unchosen actions will be (see **Figure 11C**, for an illustration).
(1) The “counterfactual (CF) learning rate” is a proxy for how much weight is given to the counterfactual prediction error. In a world where instrumental control is assumed (i.e., a world where factual and counterfactual actions give rise to different outcomes), a CF learning rate is a measure of how fast the *divergence* between factual and counterfactual outcomes builds up over time.

### CF model: best-fitting parameters

We first compared the value of the reference parameter against the “true” reference, i.e., the true mean of the reward distributions, in both experiments. The value of the fitted reference overall approximated the mean of the reward distributions (t-tests against the mean of the reward distributions in each experiment: all *p’s* > 0.05, except for the variance condition: *p* = 0.035, see **Figure 11**). Note that the value of the fitted reference varied across subjects with some participants substantially underestimating the true mean of the current distribution (see **Figure 11**, vertical dashed black lines). Interestingly, participants who underestimated the true mean also tended to exaggerate instrumental divergence as a result – i.e., the difference between chosen (factual reward) and unchosen (counterfactual reward) alternatives (see **Figure 11A and 11B**, dark solid curve, and **11C**, “True vs. Fitted Reference”).

**Figure 11.**
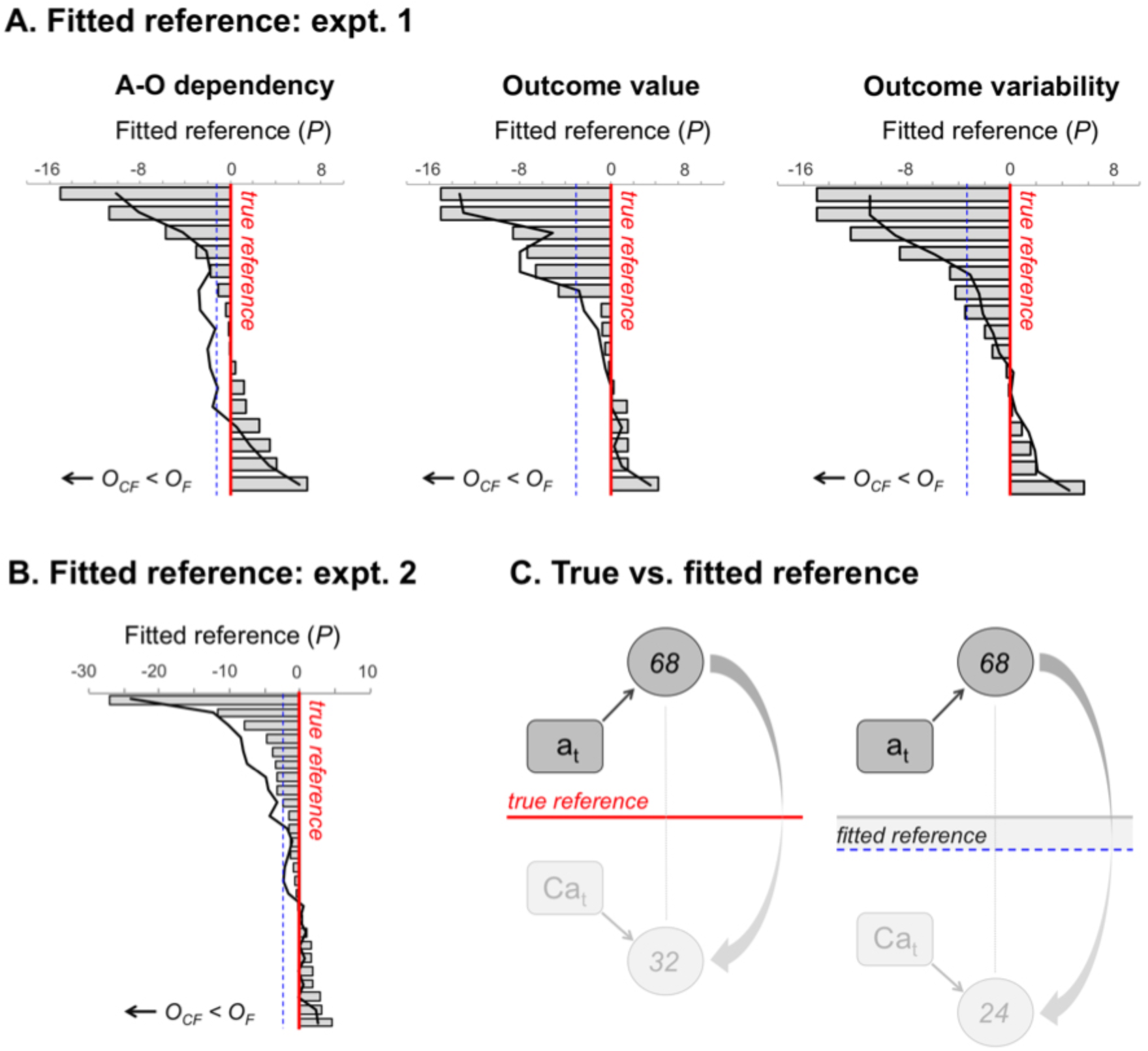
Fitted individual references across the different sessions of Experiment 1 (A) and Experiment 2 (B). The bars represent the value of the fitted reference *relative* to the true reference, i.e., the true mean of the reward distributions (vertical red line), in each participant. A negative value indicates that the participant underestimated the true reference. The greater the negative value, the lower the counterfactual reward inferred by the subject, relative to the factual reward (*O_CF_* < *O_F_*). Conversely, the greater the positive value, the greater the counterfactual reward inferred by the subject, relative to the factual one. Below the red line, the vertical dashed blue line represents the group mean of the fitted reference. Over individual bars, the solid dark curve represents the *divergence* between chosen and unchosen alternatives in each subject. The divergence was calculated by subtracting the factual from the counterfactual reward in each trial, based on the subject’s fitted reference, and averaging the result over all trials (**C) True vs. fitted reference**. When combined with the contextual rule of the CF model, underestimating the true reference leads to exaggerating the divergence between factual and counterfactual outcomes (e.g., 24 rather than 32).

We next compared factual and counterfactual learning rates within and between experiments. Three counterfactual alternatives were updated on each trial:

(i) the unchosen button of the chosen machine (α*_CF1_*),
(ii) the chosen button of the unchosen machine (α*_CF2_*),
(iii) the unchosen button of the unchosen machine (α*_CF3_*)

To first compare the factual and counterfactual learning rates of experiment 1, we carried out a 2 × 2 × 3 repeated-measures ANOVA with the button (chosen vs. unchosen), the machine (chosen vs. unchosen), and the 3 different statistics (dependency vs. value vs. variance) as within-subject factors. A similar 2 × 2 repeated-measures ANOVA was performed on all pooled conditions of experiment 2.

In experiment 1, the main effect of the “machine” (F(1,15) = 4.66, *p* < 0.03, 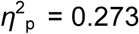) and the main effect of the “button” (F(1,15) = 15.78, *p* = 0.005, 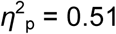) were significant. Thus, the learning rate associated with the chosen machine was significantly lower than the learning rate associated with the unchosen machine (post hoc test, all *p’s* < 0.04), whereas the learning rate associated with the chosen button was globally higher compared to the unchosen button (all *p*’s < 0.001).

The machine-by-button interaction effect was also significant (F(1,15) = 60.07, *p* < 0.001, 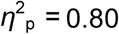). Across all three sessions of experiment 1, post hoc tests showed that learning rates of the *chosen* buttons did not differ across chosen (α_F_) and unchosen (α_CF2_) machines, while learning rates associated with the *unchosen* buttons (α_CF1_ vs. α_CF3_) differed significantly (all *p’s* < 0.001). Chosen (α_F_) and unchosen (α_CF1_) buttons of the chosen machine also differed significantly (all *p’s* < 0.001), whereas they could not be distinguished for the unchosen machine (α_CF2_ vs. α_CF3_) (**Figure 12A**). This interaction effect was observed in all conditions equally (i.e., no significant modulation of the machine-by-button interaction by the type of statistics: F(1,15) = 1.17, *p =* 0.3, 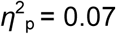). Experiment 2 showed the same tendency as observed in the previous experiment (see **Figure 12B**). However, only the machine-by-button interaction effect was statistically significant (F(1,25) = 4.06, *p* = 0.04, 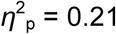). Note that the pattern of fitted learning rates from the CF model was correctly recovered when applying the procedure to simulated data, and therefore, it was not an artefact of the parameter optimization procedure (see **Figure 12C**, and **Appendix D**, “Parameter recovery procedure”).

**Figure 12.**
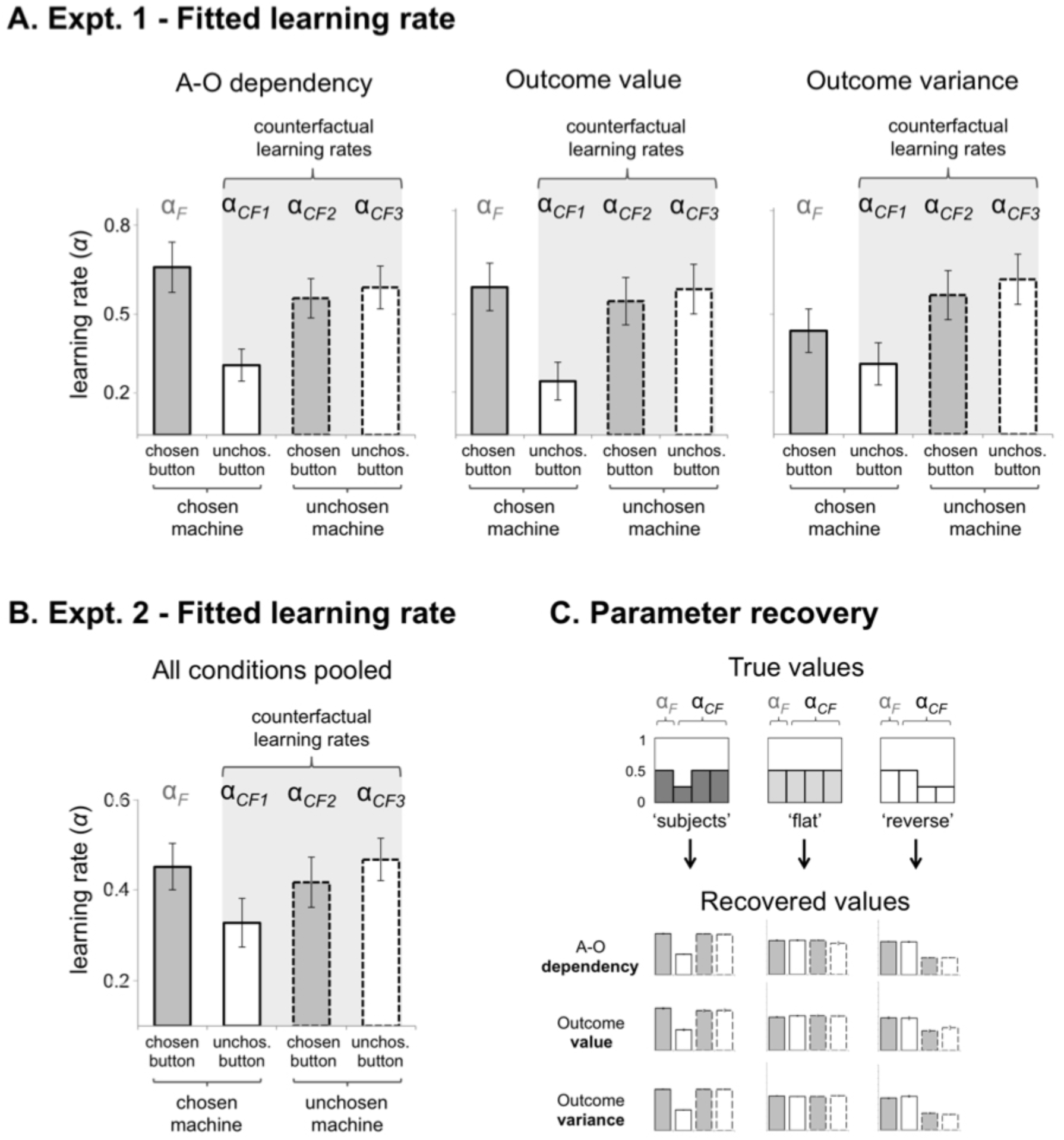
Fitted learning rates from the winning model (CF). **(A) Expt. 1:** Factual (α*_F_*) and counterfactual learning rates (α*_CF_*) within each experimental session, for each button (chosen, unchosen) and each machine (chosen, unchosen). **(B) Expt. 2**: Factual and counterfactual learning rates for all conditions pooled together. **(C) Parameter recovery procedure:** “True value”: learning rates used to simulate the data (see Appendix D, Table S1). “Recovered value”: learning rates obtained from fitting the model on the simulated data. “Subjects” = highest learning rate for the unchosen button; “flat” = identical learning rates across the unchosen button and the unchosen machine; “reverse” = highest learning rates for the unchosen machine. Our parameter optimization procedure was able to correctly recover the (true) parameter values from all patterns in all sessions.

Interestingly, our findings reveal that participants calibrate their learning rates in a way that reflects their belief about the task structure. First, counterfactual (CF) learning rates associated with the button or with the machine were significantly higher than zero in all experiments and conditions (see **Figure 12**, comparing α_CF1_, α_CF2_, α_CF3_ > 0, all *p’s* < 0.005). A high CF learning rate indicates that participants update the value of the forgone alternative; this counterfactual update results in making the value of the unchosen alternative *diverge* from the value of what is currently chosen. Thus, above-zero CF learning rates show that our participants construed their actual choice as being *causally efficient*, i.e., as making a difference relative to the unchosen alternative.

A CF learning rate is formally equivalent to the notion of “mutability” in previous work on counterfactual reasoning (e.g., Dehghani, Iliev and Kaufmann, 2012; Kahneman and Miller, 1986). Mutability is a property of a variable that signals whether the variable is likely to take different values in the real and counterfactual worlds. Thus, a *highly mutable* variable is highly likely to diverge across factual and counterfactual worlds (Lucas and Kemp, 2015). Similarly, a machine associated with a *high* CF learning rate is a *highly mutable* machine: choosing this machine, rather than the other one, should make a significant difference with respect to the outcome. Conversely, a low CF learning rate would minimize the divergence, while a *null* CF learning rate would signal a *null* divergence between the chosen and unchosen options. Importantly, our results revealed a hierarchy across buttons and machines. Counterfactual learning rates were higher for the machine than for the button, which suggests that participants conceived the former as being more “mutable” than the latter. In other words, participants believed that making a choice about the machine was more likely to make a difference to the world relative to making a choice about the button.

What does this hierarchy account for? We suggest that counterfactual emulation is more likely to be leveraged for testing control at most abstract levels of action representation (e.g., at the level of the machines) and is less required for less abstract levels (e.g., the level of the buttons) where *direct* instrumental testing is available to the subject. Crucially, should this prediction be correct, counterfactual emulation would be optimal in an environment where instrumental divergence is maximal between *machines*, rather than between buttons. We directly tested this hypothesis by simulating our CF model across two different environments: (1) an environment where divergence was maximal between buttons, or (2) an environment where divergence was maximal between machines (see **Figure 13A, left** and **right panels**, respectively, and **Appendix D**). We tested the performance of different patterns of learning rates across these two types of environment: *i*) a pattern that was similar to that of participants (*higher* CF learning rates for the most abstract level, i.e., the unchosen machine), *ii*) the reverse pattern (*lower* CF learning rates for the unchosen machine), and *iii*) a flat pattern (*equal* learning rates across unchosen machines and buttons).

Consistent with our prediction, we found that the pattern of participants outperformed the two alternative patterns in the environment where the divergence was set at the most abstract level, i.e., the machine level (see **Figure 13A**). Importantly, this result was all the more true when values of the reference point approximated the true mean of outcome distributions (see **Figure 13B**).

**Figure 13.**
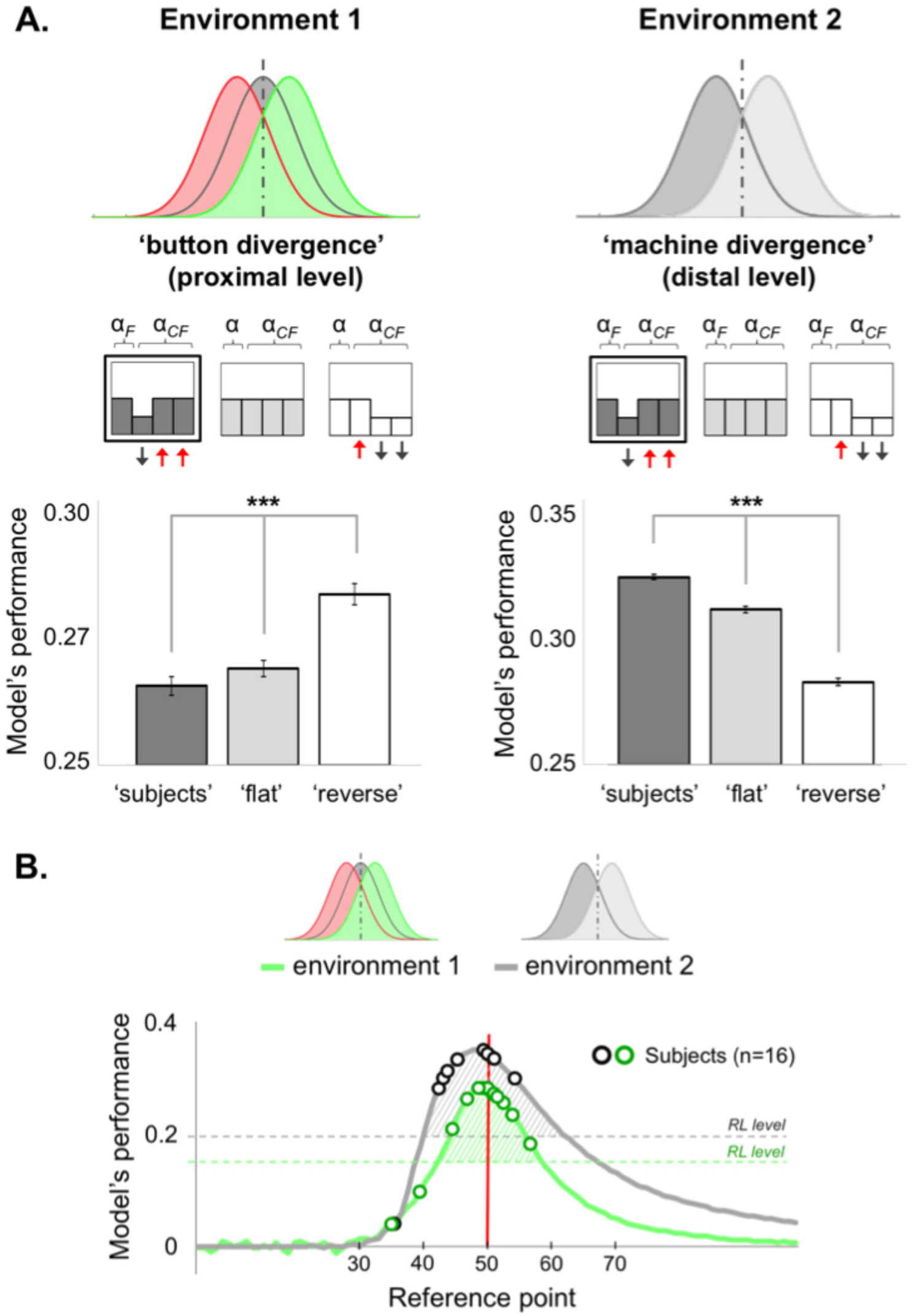
Simulated performance of the winning model (CF) across two different environments, with three distinct patterns of learning rate. **(A)** *Left panel:* performance of the CF model when the divergence is maximal between buttons (red and green distributions). The pattern of participants (dark box: “subjects”) is outperformed by the two alternative patterns (“flat” and “reverse”). *Right panel:* performance of the CF model when the divergence is maximal between machines (dark and light grey distributions). Here, the pattern of participants outperforms the two alternative patterns. Model’s performance is normalized against chance-level. α*_F_* = “factual” learning rate (chosen button of the chosen machine); α*_CF_* = “counterfactual” learning rate (unchosen button and/or unchosen machine). Three-stars: *p* < 0.001. **(B) Averaged performance of the CF winning model (*y*-axis) across the two environments, depending on the value of the reference point (*x*-axis)**. Model’s performance is optimal in both environments when the reference point approximates the true mean (red vertical line), as most subjects did (green and dark circles). The hatched areas delineate the range of reference values for which the CF model outperforms the RL model (horizontal dashed lines). In both environments, model’s performance is normalized against chance-level.

## General discussion

Using a modified reversal-learning procedure, we tested whether, and how well, human participants could adjust to self-vs. externally generated changes in a task where the source of these changes was uncertain. Specifically, any perceived changes could potentially be ascribed to three different causes: (i) the participant’s choice, (ii) the intrinsic variability of the outcome, or (ii) a reversal in either instrumental or environmental contingencies. Thus, maximizing performance in the task required the ability to discriminate action-related changes from changes due to intrinsic feedback noise and/or external volatility as well as to adjust one’s choice behaviours accordingly.

## Behavioural results: control, value and variance

In all experiments, we found that participants were able to discriminate the best-from least-rewarding buttons and to distinguish between the controlled and non-controlled machines – that is, between the machine for which there was a best- and a least-rewarding button and the machine for which both buttons were equally rewarding. Participants performed the task well despite the lack of explicit cues to signal reversals in the best-rewarding button or in the controlled machine. In experiment 1, participants preferentially chose the controlled machine over the non-controlled machine while also exhibiting a marked preference for the best-valued and low-variable machines in both experiments 1 and 2. Interestingly, both outcome value and variance had an effect on the proportion of the *controlled* machine. Thus, participants more often chose the controlled machine when the relative *value* of this machine increased but also when its relative *variance* decreased. Likewise, when the value of control increased, participants discriminated better between the best and worst options of the controlled machine, and choice behaviour was therefore found to better reflect the true divergence of the controlled machine.

This interaction between value and control, as well as between variance and control, is reminiscent of self-attribution biases in which healthy adults take credit for positive outcomes while denying responsibility for negative events (e.g., Mezulis *et al*., 2004), and overestimate the variability of random series while under-estimating the variability of self-caused events (e.g., Boland and Pawitan, 1999). Spurious *positive* relationships between control and value are further exemplified in situations where people mistake the value of an event for real control over this event through inflating probabilistic estimates of action-event contingencies (Kool, Gatez and Botvinick, 2013). This interplay between control, (high) reward value, and (low) variance, suggests that individuals construe the effects of their action along multiple dimensions: as a mean to make a difference to the world but also as an instrument to bring about positive events and to reduce the inherent variability of the environment. Importantly, we found that one single model (CF) was able to simulate participants’ performance across all three dimensions and was able to do so with both the same set of parameters and same parameter values.

## Associative learning and counterfactual update

In all experiments, optimal performance required a complete knowledge of the underlying causal structure of the task, i.e., a representation of each probability distribution relating each possible action to a particular state. Thus, reaching optimal performance was computationally costly because it ideally required maintaining probability distributions across all alternative causes and updating all possible alternatives at once whenever integrating new evidence. Whether such a strategy is used, or is even usable, by human subjects remains unknown (Eckstein *et al*., 2004; Jones and Love, 2011). Although they lack an explicit representation of instrumental contingencies, simpler learning schemes, e.g., based on pure associative processes, can readily adapt to causally structured environments, at a much lower cost (Dickinson, 2001). On the other hand, associative processes only enable a form of *proximal* instrumentality in which acquisition and performance of new and existing behavioural strategies are regulated by their immediate consequences. Accordingly, associative agents only slowly adapt to environments with periodically changing action-outcome mappings and therefore would hardly approximate the efficiency of human performance (Gershman, Markman and Otto, 2014). An intermediate solution would consist of combining a (simple) associative learning scheme with a generative rule for emulating an approximate version of the environment’s causal structure. In contrast with pure associative algorithms, this “combined” model would assume a generative source behind observation, but this source would not have to be a fully specified probability distribution of expected action outcomes. Models of *counterfactual reasoning* (e.g., Lucas and Kemp, 2015) can be specified to permit action outcomes to take different values in the real and counterfactual worlds. Importantly, these models can also account for hierarchical inference in causal reasoning by allowing factual and counterfactual action values to be updated according to different dynamics (e.g., learning rates).

We tested and compared the ability of associative, generative, and counterfactual models to account for the participants’ data across all experiments. We found that models that *merely* aimed at maximizing action value –whether by prediction-error minimization (RL) or by means of Bayesian inference (BM)– could not explain the participants’ choices well and neither could a model (BC) that aimed to maximize control over the task, regardless of action value. For both quantitative (BIC) and qualitative (reversal curves) criteria, participants’ behaviour was best accounted for by a model that made choices based on counterfactual contingencies, i.e., a model that emulated *unseen* action-outcome pairs according to a contextual rule. Thus, counterfactual contingencies were emulated by deriving the consequences of the forgone action from the current action taken and by assuming relative (i.e., context-dependent) *divergence* between both. Importantly, instrumental divergence was implemented in the model as a prior.

Specifically, this prior conveys the belief that taking a specific action (e.g., choosing option A vs. B) makes a difference in terms of the outcome. As mentioned above, instrumental divergence is a reliable proxy for goal-directed control because the more the action *diverges* with respect to its contingent states (the factual and counterfactual outcomes), the more flexible control one has over the environment. The fact that the CF model best explained the data in all conditions suggests that human subjects construe their causal power based on such a prior. Interestingly, the CF model also provides the best match for the participants’ choice even when no true control existed (i.e., “value” and “variance” conditions), which suggests that this prior could be considered a default belief in which goal-directed actions are thought to be *causally efficient* (i.e., divergent) in nature. Finally, only the CF model was able to replicate the value-by-variance interaction observed in subjects for both machine and button choices, while the other models replicated this pattern only partially (e.g., RL) or exhibited the reverse performance pattern (e.g., BC) (see **Figure 10**).

## The counterfactual world negatively covaries with the real world

Counterfactual reasoning has been the subject of many investigations in the decision-making domain from regret-based theory of choice (e.g., Bell, 1982; Coricelli *et al*., 2005) to empirical works on fictive learning, i.e., learning from alternative action values (e.g., Lohrenz *et al*., 2007). While it has been repeatedly shown that instrumental learning benefits from tracking alternative courses of action and their outcomes, how these counterfactuals are generated, and based on what rule, is currently unclear. In most studies on fictive learning, subjects are explicitly informed about the result of the forgone alternative (e.g., Lohrenz *et al*., 2007; Boorman, Behrens and Rushworth, 2011; Palminteri *et al*., 2015). In our task, however, the reward associated with the unchosen machine was not shown to the participant but had to be inferred given the chosen button. Crucially, our CF model provides an algorithmic explanation for how counterfactual action values were inferred according to a flexible, context-dependent reference, whose value was separately fitted to each participant’s data.

As previously argued, exploiting counterfactual information can be beneficial to the learner if the cost of getting and storing the information is not too high (Boorman, Behrens and Rushworth, 2011). Importantly, the context-dependent reference embedded in the CF model approximated the *mean* of the generative distributions in the task, and thus allowed for emulating counterfactuals to occur with a low cost. The mean is a simple and often-optimal operator that minimizes prediction error. Under uncertainty, making decisions based on an averaged representation of the environment is often advantageous (Sutton and Barto, 1998). In this respect, the CF model would be efficient, not because it would maintain an expensive, yet accurate, causal model of the task (e.g., the probability distributions over all possibilities), but because it embeds a prior (the reference point) that approximates the actual structure of the environment (see also Parpart *et al*., 2017). In addition to showing better performance than a classical RL, the CF model also maintains simplicity in terms of algorithmic design and computation. We argue that this simplicity provides a step towards an explanation of how human agents achieve a trade-off between robust causal inference and the costs of maintaining an accurate model of the world (Bramley *et al*., 2017).

Updating the counterfactual according to a context-dependent reference is consistent with a broader literature on reference dependence in behavioural economics in which the utilities of outcomes are assessed relative to a context-specific reference point (e.g., Kahneman and Tversky, 1979; Köszegi and Rabin, 2006; Denrell, 2015). Converging evidence from reinforcement comparison methods and behavioural economics equally suggests that people make decisions according to a context-dependent reference (Palminteri *et al*., 2015; Palminteri *et al*., 2017; Klein, Ullsperger, & Jocham, 2017; Burke *et al*., 2016). Importantly, providing counterfactual information to the subject reinforces the dependence on context for evaluating rewards and punishments (Palminteri *et al*., 2015). Thus, when subjects are informed about the result of the forgone alternative, value contextualization is enhanced. Similar to our CF model, such contextualization would consist in tracking the mean of the distribution of values of the current choice context (i.e., the reference point) and using it to centre both factual and counterfactual option values. Such value contextualization echoes adaptive coding of outcomes in neural populations in which neural outputs rescale to the range of currently expected outcomes (Burke *et al*., 2016), and it is more generally consistent with studies on context-based processing of outcome information showing that motivationally relevant information is encoded in a relative fashion that adapts to the current value-context (Seymour and McClure, 2008).

Because it updates alternative action values based on a context-dependent reference, the CF model can be viewed as a generalization of the Rescorla-Wagner (R-W) rule (Sutton and Barto, 1998). Interestingly, counterfactual updating in associative learning can also be modelled using a Bayesian generalization of the R-W model, i.e., using Kalman filters based on temporal difference (TD) learning (Keramati, Dezfouli and Piray, 2011). Kalman TD incorporates a component of counterfactual thinking by encoding a *negative covariance* between stimuli elements. In terms of instrumental learning, this covariance structure can be leveraged to update both factual and counterfactual action values, as learning one particular instrumental contingency automatically leads to a reduction in the associative strength of the unchosen contingency (Gershman, 2015). In a recent study, Morris *et al*. (2017) showed that instrumental learning was best explained by a Kalman algorithm that combines prediction-error learning with a similar covariance matrix, which reflects the structure of the task environment. In their task, several causal variables compete to explain the observation. The winning model assumes negative covariance between these variables, which means that a change in the belief of one cause inversely affects the other (Morris *et al*., 2017). The covariance matrix thus allows the learner to reason *counterfactually* about alternative courses of action, which allows her differentiate the unique effects of action from background effects, i.e., from effects that would have occurred in the absence of that action. Morris and collaborators found that this model, which combines key features of associative learning and model-based RL, better characterized behavioural performance and neural activity associated with instrumental learning than models based on covariance or prediction error alone.

Morris *et al*.’s model has formal resemblance to our CF model. In the CF model, however, the negative covariation between factual and counterfactual values critically relied on a parameter, the reference point, whose value was separately fitted in each participant. Importantly, the value of this reference showed some variability across participants, depending on their subjective preferences and beliefs. Thus, while some underestimated, some other overestimated the true mean of the current distributions. Interestingly, underestimating the true mean was *self-serving* in nature because it led participants to exaggerate the divergence between factual and counterfactual outcomes. Thus, in subjects underestimating the true mean, the lower the reference, the worse the outcome would have been had they made *another* choice (*O_CF_* < *O_F_*, **Figure 12**). Conversely, participants overestimating the true mean could be seen as pessimistic because they assumed that the alternative course of action would have been better off on average (*O_CF_* > *O_F_*, **Figure 12**). This result agrees with a variety of empirical works showing that while healthy adults exhibit attribution biases when judging their agency, these biases vary substantially across individuals (Mezulis *et al*., 2004, for a review). By combining counterfactual updating with a subjective reference point, the CF model accounts for interindividual variability in self-serving beliefs, i.e., perceived controllability, during online instrumental learning.

## Counterfactual emulation operates at most abstract levels of action control

Negative covariance is at the heart of the notion of “difference-making” in counterfactual theories of causal reasoning. Counterfactual (CF) theories posit that a cause is something that makes a difference to another event (Walsh and Sloman, 2011). According to the CF view, individuals would infer causal relations by simulating models of close alternatives (“nearest possible worlds”) in which the candidate cause (A) is negated and the outcome is observed (O). If the outcome is undone (~O) as a result of negating the candidate cause (~A), then the probability that A is selected as the cause should increase accordingly (e.g., Roese & Olson, 1997; Woodward, 2003; Sloman and Lagnado, 2015). When applied to intentional causation, an action should be considered “causal” when simulating a change in that action (e.g., the action is not taken) produces a change to the outcome (e.g., the outcome does not occur). In CF theories, “mutability” is a property that characterizes the effects of “simulating” changes in one variable, and hence can be seen as a measure of its causal power (Kahneman and Miller, 1986; Dehghani, Iliev and Kaufmann, 2012). Thus, a variable is highly mutable if realizing, relative to not realizing, this variable is likely to make a difference in the world. Put differently: a variable is mutable if it is likely to *diverge* across factual and counterfactual worlds (Lucas and Kemp, 2015).

The notion of “mutability” closely relates to the notion of counterfactual learning as instantiated in the CF model through a *counterfactual (CF) learning rate* parameter. A CF learning rate can be viewed as measuring the *speed of divergence* between factual and counterfactual outcomes. Thus, the greater the value of the CF learning rate, the faster the counterfactual action value is assumed to diverge from the chosen action value. In our task, a machine associated with a *high* CF learning rate is a *highly mutable* machine: choosing this machine, rather than the other one, is thought to make a significant difference with respect to the outcome. Importantly, our results revealed that counterfactual learning was hierarchically organized: CF learning rates were higher for the *machine* than for the button, which suggests that participants conceived the former as being more “mutable” than the latter. In other words, participants considered that making a choice about the machine was more likely to make a difference to the world relative to making a choice about the button.

This hierarchy in learning from factual and counterfactual action values might be well adapted to an environment where changes in causal influence (e.g., reversals) operate at more or less abstract levels of action control. Higher learning rates for the CF machine relative to the CF button suggest that subjects are more likely to engage in counterfactual emulation for testing their control at most abstract levels of action representation (the machine), and less for less abstract levels (the button) where *direct* instrumental testing is available. Should this prediction be correct, such a hierarchy in counterfactual learning would be advantageous in an environment where instrumental divergence is maximal between *machines* rather than between buttons. We directly tested this hypothesis by simulating our CF model across two different environments, where maximal instrumental divergence was either between buttons or between machines. In line with our predictions, we found that the CF model was best suited to deal with an environment where the divergence was set at the most abstract level, i.e., the machine level (see **Figure 13A**).

That individuals are more likely to engage in counterfactual emulation for the most abstract levels of action control is consistent with evidence from hierarchical models of action representation (e.g., Kilner, 2011; Chambon *et al*., 2017). In such models, an observer predicts another person’s behaviour based on beliefs derived from simulating one’s internal model (i.e., a model of how people are likely to behave in a given context). The nature of these beliefs critically depends on the level at which the behaviour is represented, from the least to the most abstract levels (e.g., kinematic vs. motor vs. goal level). Thus, a change at the most abstract level (e.g., the goal level: going to restaurant *vs*. theatre) is assumed to have a greater effect on the resulting behaviour than a change made at a *less* abstract level (e.g., the kinematic level: using a power vs. precision grip to grasp a mug). Importantly, human subjects show greater reliance on their internal models when having to predict more and more abstract behaviours (e.g., going to the restaurant vs. theatre > using a power vs. precision grip) (Chambon *et al*., 2011; Chambon *et al*., 2017). Likewise, our results indicate that human subjects are more likely to emulate counterfactual alternatives when making decisions at more abstract levels of action control (machine > button).

## Control beliefs foster opportunities for learning

Assuming a *negative* covariance between factual and counterfactual outcomes implies that the world can be divided into states that are essentially anti-correlated. In this scenario, only two states are possible: you are the agent or you are not. This assumption agrees with the fact that judgements of agency are *binary* in nature. Indeed, while individuals readily experience intermediate levels of sensorimotor control, confidence, or difficulty, they rarely, if ever, experience *intermediate* levels of agency; they can be “more or less confident”, but they do not feel “more or less agent” (Chambon and Haggard, 2013). The all-or-none nature of agency is supported further by the observation that people think of causal relationships between actions and outcomes in terms of “state” (is A the cause of O?) rather than in terms of “force” (Tenenbaum and Griffiths, 2001). We show that the CF model best fits participants’ behaviour because it embeds a prior that matches one’s belief about the structure of controllable environments in which binary and abrupt, rather than smooth and continuous, changes in contingencies can occur. In this sense, the CF model would be best suited to track changes in agency (me vs. not-me) than gradual changes due to external volatility (e.g., the light decreasing over the course of a day).

We speculate that this prior about negative covariance mirrors participants’ beliefs about their control over machine outcomes: had their choice been different, the outcomes would also have been different. Importantly, participants hold this control belief even in sessions where no true control existed (see Experiment 1, “value” and “variance” sessions), or although instructions made no reference to control in the task (see Experiment 1b, **Supplementary information**). Beliefs in one’s causal power are a strong determinant of voluntary behaviours: individuals are more likely to enact certain behaviours when they feel or believe they can enact these behaviours successfully (e.g., Ajzen, 2002). Control beliefs develop early and are somewhat irrepressible: the need to be and feel in control is so strong that individuals would do whatever they can to re-establish control when it disappears or is taken away, including self-attributing unrelated events (Langer, 1975) or acting superstitiously in the belief that their action is accountable for uncontrollable outcomes (Blanco, Matute and Vadillo, 2011). Importantly, control beliefs would explain an enduring puzzle in causal reasoning, that is, why people show remarkable performance in causal inferences, which they often make effortlessly and from very little data, and yet readily experience illusory control, whether in real-life uncontrollable situations (Matute, 1996) or in experimental settings with null contingency (Blanco, Matute and Vadillo, 2011). This relationship between illusory control and control beliefs is further corroborated by people’s tendency to self-attribute positive outcomes when their perceived controllability of the environment is high (Harris and Osman, 2012).

One explanation for assuming control as a default belief – whether illusory or not – is *learning*. Indeed, control beliefs would be particularly adapted to controllable environments, whose latent causal structure can be *learnt* to maximize rewards in the long run (Lake *et al*., 2016). In a structured environment, enacting actions, relative to not acting, is advantageous on average as the reward/punishment ratio can be turned in favour of rewards though implementing appropriate actions. In this situation, agents would be better off holding the belief that their actions are efficient means for attaining desired outcomes. In sum, the detrimental consequences of assuming control as a default belief would be offset by opportunities for learning the causal structure of the world, and therefore by the ability to flexibly switch preferences when reversals occur, which ultimately reduced the cost associated with missing opportunities (Koechlin, 2016). Importantly, control beliefs, such as self-efficacy, play a major role in general health and wellbeing (Bobak *et al*., 2000). A lowered sense of causation is associated with lower self-esteem, greater mood disorders and greater depressive symptoms (Bandura, 1997). Depressed individuals perceive their environment as being more random than non-depressed people. In the depressed view (so-called “depressive realism”), the reward/punishment ratio is evenly balanced, which substantially reduces opportunities for learning and ultimately makes any action pointless (Nettle and Bateson, 2012). This account agrees with clinical reports of greater passivity in depressed people, that is, a reduced ability to initiate voluntary actions (Blanco, Matute, & Vadillo, 2012). Acting with less frequency would make depressed individuals exposed to fewer *incidental* associations between actions and action-contingent events (reduced “action-density” bias, see Matute *et al*., 2015), which might in turn impede learning instrumental contingencies and aggravate depressive symptoms in the long run.

The strength of the CF model stems from the simplicity with which it embeds the participant’s prior beliefs about control. This prior amounts to assuming relative (i.e., context-dependent) divergence between factual and counterfactual worlds. We argue that this simple prior allows the model to rapidly and flexibly switch preferences when a reversal occurs, as demonstrated by its robust learning curves and performances in both experiments, relative to more sophisticated models such as those aiming at statistical optimality (e.g., the BC model). We believe that simplicity is required to account for the ease with which resource-bounded agents learn instrumental contingencies but also to explain how strong control beliefs can be sustained as a default backdrop to our normal mental life. As mentioned above, a pervasive belief in one’s causal power can make instrumental learning sometimes depart from statistical optimality, which results in illusions of control and an inflated perception of one’s own efficacy. The influence of such a pervasive belief would explain why learning action-outcome causal relationships seems not to suffer the same biases as other forms of causal learning that are based on passive observation (Morris *et al*., 2017; Chambers *et al*., 2017). Ultimately, strong control beliefs in human agents could account for why reasoning about *external* causes differs from reflecting upon one’s own causal power both in terms of underlying computations, normative principles and optimal behaviour.

## Conclusions

We designed a series of experiments that required participants to continuously monitor their causal influence over the task through discriminating changes that were caused by their own actions from changes that were not. Comparing different models of choice, we found that participants’ behaviour was best explained by a model (CF) deriving the consequences of the forgone action from the current action taken and assuming relative divergence between both. Importantly, this model agrees with the intuitive way of construing causation as “difference-making” and further endorses the long-held view that goal-directed actions are *divergent* in nature: they make a difference to the world and can hence be implemented as efficient means for pursuing desirable outcomes. In the CF model, difference-making was explicitly accounted for by assuming *negative covariance* between factual and counterfactual action values. Based on this prior, the CF model directly *emulated* counterfactual action values through a subjective reference point that aligned with the actual structure of the task environment. Crucially, we found that counterfactual emulation was more likely to occur at most abstract levels of action control, in line with evidence from hierarchical models of goal-directed actions. Finally, we suggest that the CF model outperformed all competitors because it closely mirrors people’s beliefs in their causal power, which are beliefs that are well suited to learning action-outcome associations in controllable environments. We speculate that control beliefs may be part of the reason why reflecting upon one’s own causal power fundamentally differs from reasoning about external causes.

## Acknowledgments

We thank Valentin Wyart and Stefano Palminteri for comments and useful discussions about earlier versions of this manuscript. V.C. was supported by the Agence Nationale pour la Recherche (ANR) grants ANR-17-EURE-0017 (Frontiers in Cognition), ANR-10-IDEX-0001–02 PSL* (program “Investissements d’Avenir”), and ANR-16-CE37–0012–01. H.T. was supported by a PSL/ENS studentship. E.K. was supported by a European Research Council Grant (ERC-2009-AdG #250106). C.F. was supported by a DGA/INSERM fellowship.

## Appendix A

### Generative model

As mentioned in the main text, the generative model was a Bayesian learner that updated beliefs associated with each possible action on each trial. Specifically, the model (either BM or BC) aims to infer the correct action mapping between the four possible mappings (or states). We further define a state-function *f* specifying the underlying reward distribution for a given action *a*_1,…,4_ and state *z*, as follows:

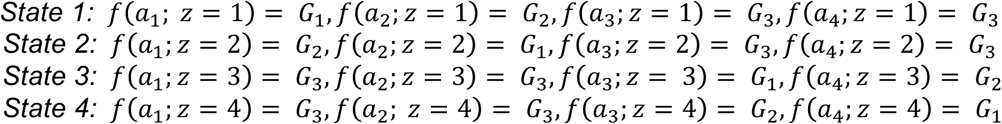

where *a*_1,…,4_ corresponds to each possible action (among the 4 possible combinations of button and machine), *G*_1_ is the distribution associated with having selected the best-rewarding button of the controlled machine, *G*_2_ is the distribution associated with having selected the least-rewarding button of the controlled machine, and *G*_3_ is the distribution associated with having selected the non-controlled machine (see respectively green, red, and grey distributions of Figure 4, top panel). We assume the rewards to be drawn from Gaussian distributions, as they were in the task (see Figure 5A).

The analytical model used to infer the state on each trial is a hidden Markov model, defined as follows:

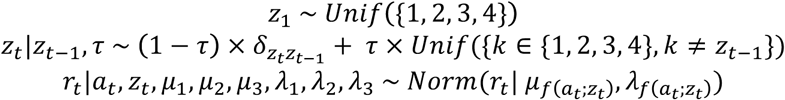

where *z_t_* corresponds to the state inferred on trial t, *r_t_* corresponds to the reward observed on trial t, *a_t_* corresponds to the action observed on trial t, and *δ* is the index function, i. e., 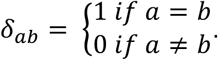. Note that the reward values *r_t_* were rescaled within the range [0,1].

Let the parameters be *θ* = (*τ*, *μ*_1_, *μ*_2_, *μ*_3_, *λ*_1_, *λ*_2_, *λ*_3_). The analytical model to infer the parameters on the first trial, given their prior hyperparameters, is the following:

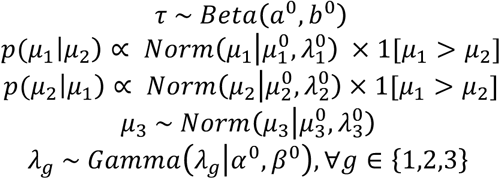

where *τ* is the volatility parameter, and *μ_g_* and *λ_g_* are the mean and precision of the Gaussian *G_g_* for *g* ∈ {1,2,3}. To obtain conjugate distributions, we used Gaussian precisions rather than their standard deviations. We assumed the same hyper-parameters for the precisions of the three Gaussians. These hyper-parameters led to the less informative priors possible and were set to the following values: ***a*^0^** = 1; ***b*^0^** = 9; 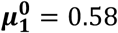; 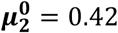; 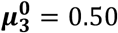; 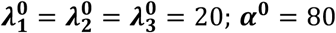; ***α*^0^** = 80; ***β*^0^** = 0.8.

**Figure S1.**
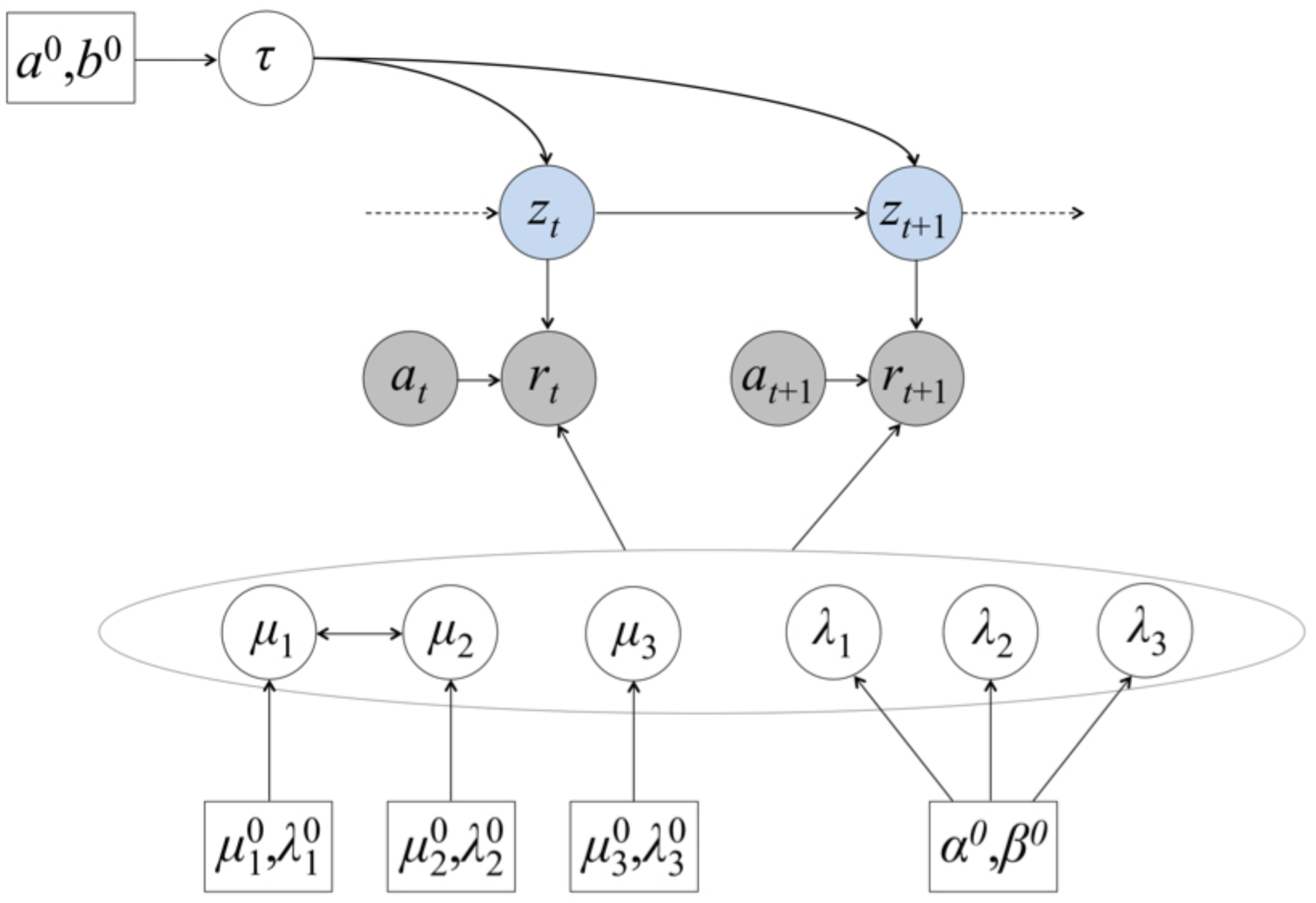
Generative graphical model. assumed by the subject. The variable *z_t_* corresponds to the state on trial *t* (shown in light blue). *r_t_* and *a_t_* are the observed variables (shown in grey): *r_t_* corresponds to the reward observed on trial *t*, and *a_t_* corresponds to the action observed on trial *t*. The parameters are *θ* = (*τ*, *μ*_1_, *μ*_2_, *μ*_3_, *λ*_1_, *λ*_2_, *λ*_3_). shown in white circles: *τ* is the volatility parameter, and *μ_g_* and *λ_g_* are the mean and the precision of the Gaussian *G_g_* for *g* ∈ {1,2,3}. The hyperparameters are shown in white boxes.

The goal of the inference algorithm is to predict the state of the next trial:

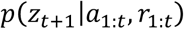

Let us have *I* = 1,000 samples 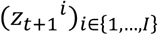 distributed under the distribution *p*(z*_t_*_+1_ |*a*_1:_*_t_*,*r*_1:_*_t_*). A Monte Carlo approximation of the integral gives:

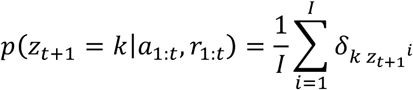

To perform inference, we used a Gibbs algorithm sampling iteratively the latent states and the parameters.

Regarding the sampling of the latent states, for the first trial, the posterior distribution on the hidden state takes the following form:

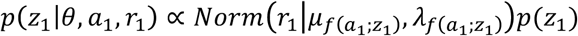

Then, the forward recursion from trial *i* − 1 to trial *i* leads to:

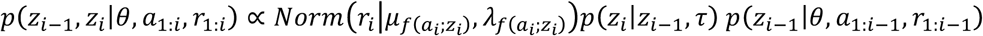

with 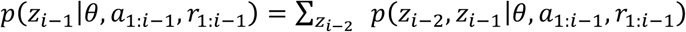

The latent sample *z*_1:_*_t_* is thus obtained by drawing *z_t_* from *p*(z*_t_*|*θ, a*_1:t_, *r*_1:_*_t_*), and then by iteratively sampling backward *z_i−_*_1_|*z_i_~ p*(., *z_i_*|*θ, a*_1:i_, *r*_1:_*_i_*) (Scott, 2002).

For the parameter sampling step, the posterior distribution of the volatility parameter *τ* depends on the number of switches predicted by the sampling trajectory *z*_1;t_. Let us define *N_switch_ = card*(*i* ∈ {2,…, *t*} | *z_i_* ≠ *z_i_*_−1_). The posterior distribution of the volatility parameter is updated as follows:

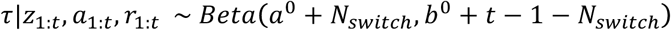

The means and precisions of the three Gaussians are also updated based on past history of actions and rewards. This update first requires identifying which Gaussian *g* each observed reward was drawn from. Let us define *I_g_* = {*i* ∈ {1,…, *t*} | *f*(*a_i_*; *_zi_*) = *G_g_*}. Then, the number of trials in which the rewards are drawn from the Gaussian *g* is: *N_g_* = *card*(*I_g_*), and the average reward observed for the Gaussian *g* is: 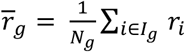.

To sample from the mean of *G*_3_, the Gaussian associated with the non-controlled machine, one just computes the tractable posterior and samples from it:

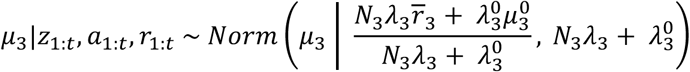

For the means of *G*_1_ and *G*_2_, the Gaussians associated with the best- and the least-rewarding button of the controlled machine, the additional inequality constraint makes the posterior distribution intractable. We thus use Monte Carlo Markov Chain procedures within the Gibbs algorithm to sample from the constrained conditional distributions of *μ*_1_ and *μ*_2_. For the proposal distribution of the Metropolis-Hastings algorithms implemented, we use the unconstrained posterior:

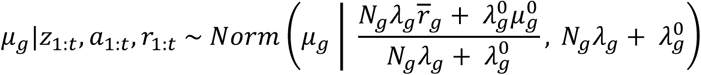

with *g* ∈ {1,2}.

As for the posterior distributions of the precision parameters *λ_g_*, they are of the form:

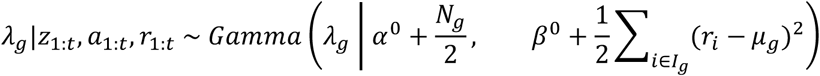

with *g* ∈ {1,2,3}.

## Appendix B

### Experiment 1b: Methods and Results

20 participants (11 females, age 21–29) took part in the experiment. The task (stimuli, timeline, and trial structure) was identical to that used in experiment 1. The only differences were the instructions and the order of sessions. The verbal and written instructions did not make any reference to a controlled machine or to a best-rewarding button. Thus, participants were only instructed to choose the option that would maximize their total reward while being reminded that they would always win the *sum* of the two machines on each trial. The order of the three experimental sessions was fully randomized so that participants could begin by any of the three sessions (dependency, value, or variability).

Experiment 1b comprised the same number of episodes as in experiment 1, with similar length and same number of button and/or machine reversals. The same four models (RL, CF, BC, BM) were fitted to participants’ choices and simulated over all three sessions with their best-fitting parameters.

The results replicated the experiment 1. Participants discriminated well between the two buttons of the controlled machine in the dependency session only (best- and least-rewarding buttons: 0.33 vs. 0.20, t(19)= 3.96, *p* < 0.001) while choosing equally button 1 and button 2 of the non-controlled machines in all sessions (all t’s < 2.16, all *p*’s > 0.12). Second, we tested whether participants showed a preference for one machine over another within each session, by comparing choice proportion for each machine against the chance level (0.5). As again expected, we found that participants showed a significant preference for the *controlled* machine in the dependency session (t(19) = 1.98, *p* < 0.05) and a marked preference for the *best-rewarding* machine in the value condition (t(19) = 5.97, *p* < 0.001). In contrast to experiment 1, however, they only showed a tendency to prefer the *low-variable* machine in the variance condition (t(19) = 1.2, *p* = 0.19).

As in the previous experiment, we compared “button” preferences across all sessions by subtracting choice for one button from choice for the other button within each preferred machine. The one-way ANOVA confirmed that “button” preferences differed across the 3 sessions (F(2,57)=12.23, *p* < 0.001, 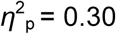). Thus, participants discriminated between buttons of the preferred machine in the *dependency* session to a greater extent than in the value and variance sessions (post hoc tests: all *p*’s < 0.001). We then compared the proportion of choice for the preferred machine across all 3 sessions. The one-way ANOVA revealed that “machine” preferences differed across the 3 sessions (F(2,57)=17.95, *p* < 0.001, 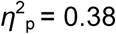), Thus, participants chose the *best-rewarding* machine (“value” session, 0.65) more than the *controlled* machine (“dependency” session, 0.54), and more than the *low-variable* machine (“variance” session, 0.52) (post hoc tests, all *p’s* < 0.001). Finally, participants were able to adjust to machine and/or button reversals, on average reaching the plateau of performance around 10 trials after reversal.

Again, replicating experiment 1, the CF model best predicted participants’ choices than all other models in the set (RL, BM or BC), in all three sessions (exceedance probability > 92%) (see **Table 1**).

### Experiment 1b: Preliminary discussion

Most of the results from the previous experiment were replicated. As expected, participants exhibited a strong preference for high rewards and preferentially chose low-variable machines, regardless of their overall value. We also found that participants were able to discriminate between causally efficient actions (the two buttons) and to identify *where* in the task environment choosing one action rather than another made a difference to the outcome (the controlled machine), and where it did not (the non-controlled machine). Importantly, this pattern of choice preference could neither be explained by the session order nor by the instructions. In this experiment, participants could as well start with the “dependency” as with the “value” or the “variability” sessions. Furthermore, participants were only instructed to make choices that maximized their rewards over both machines and time. Finally, in this experiment as in the previous one, we found that a model based on a simple context-dependent counterfactual rule (CF) outperformed all competing models, including a pure reinforcement learner (RL) and a model that explicitly aimed at maximizing reward by means of Bayesian inference (BM).

## Appendix C

### Experiment 2: Supplementary figures

**Figure S2.**
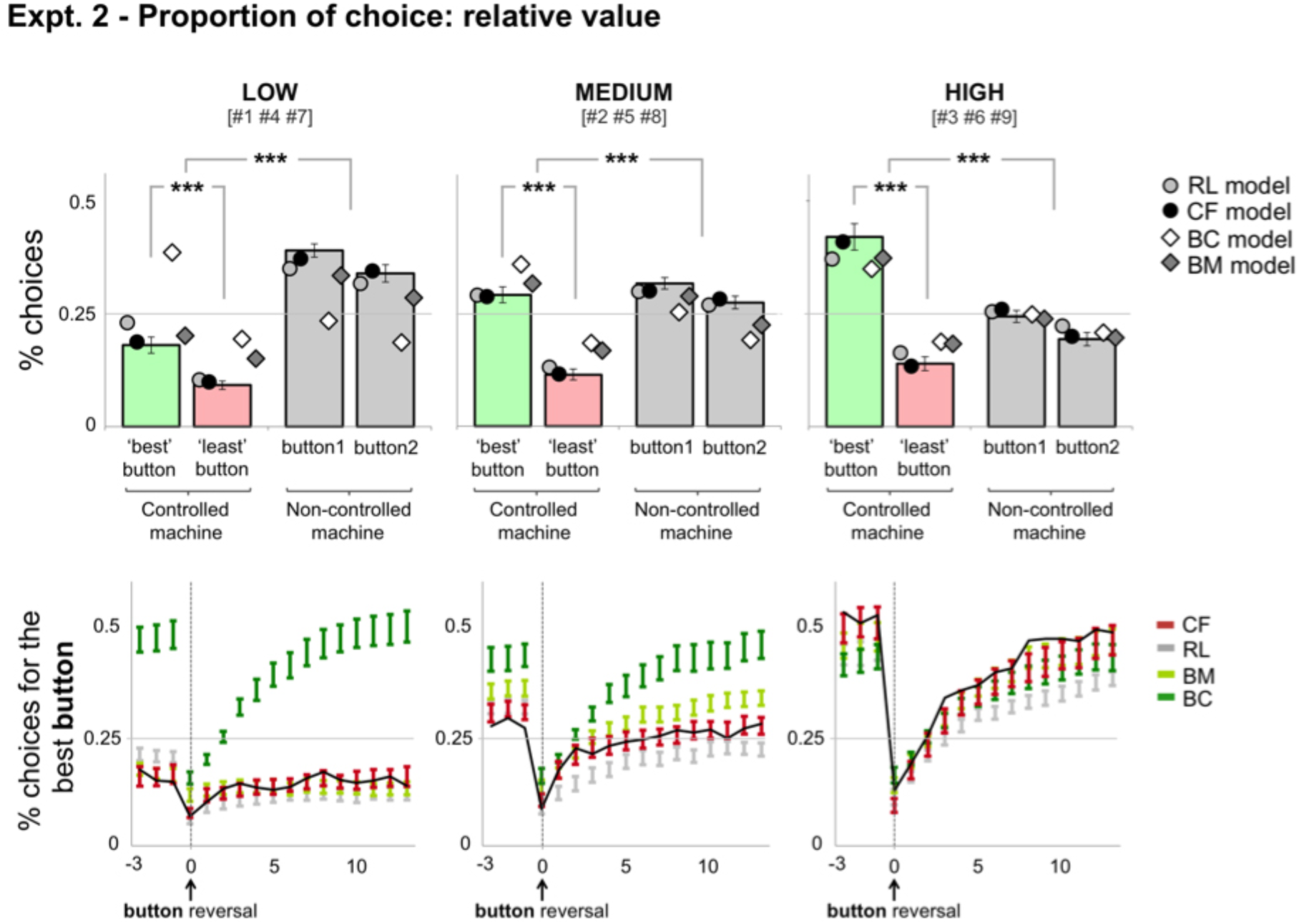
*Top panel:* Mean proportion of choice for each button of each button across the “value” dimension. Low, medium, high: the value of the controlled machine was low, medium, or high, *relative* to the value of the non-controlled machine. The numbers between brackets refer to experimental conditions shown in Figure 5. All error bars indicate standard error. For the sake of visibility, models’ error bars are not shown. Three-stars: *p* < 0.001. *Bottom panel:* **Reversal curves for participants (solid black line) and models (colored bars), across all three levels of the “value” dimension**. Model simulation: CF (red bars); RL (light grey); BM (light green bars); BC (dark green bars). Bars indicate standard error. Dashed vertical lines indicate reversal point. Machine reversal curves are not shown.

**Figure S3.**
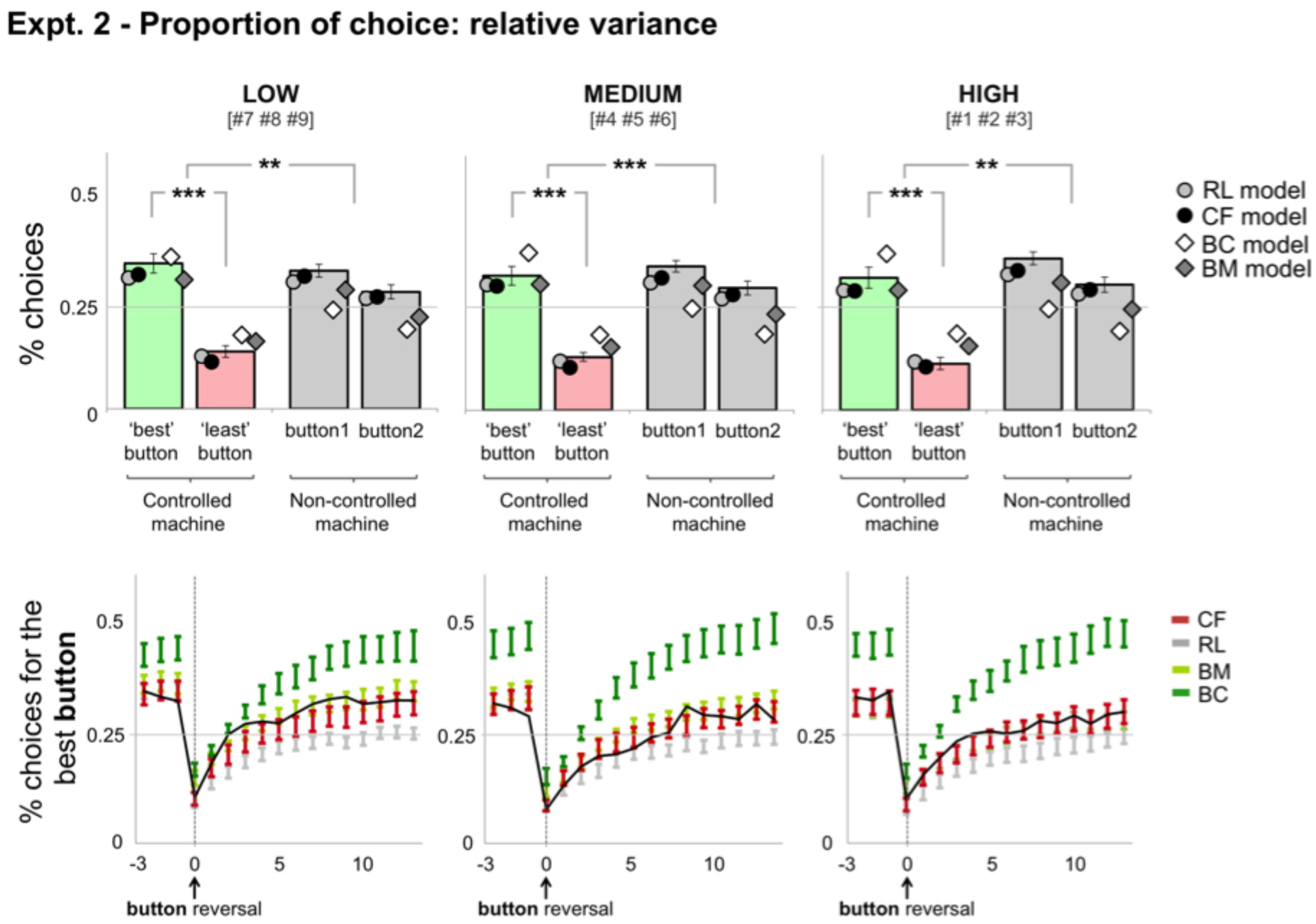
*Top panel:* Mean proportion of choice for each button of each button across the “variance” dimension. Low, medium, high: the variance of the controlled machine was low, medium, or high, *relative* to the variance of the non-controlled machine. The numbers between brackets refer to experimental conditions shown in Figure 5. All error bars indicate standard error. For the sake of visibility, models’ error bars are not shown. Two-stars: *p* < 0.01; Three-stars: *p* < 0.001. *Bottom panel:* **Reversal curves for participants (solid black line) and models (colored bars), across all three levels of the “variance” dimension**. Model simulation: CF (red bars); RL (light grey); BM (light green bars); BC (dark green bars). Bars indicate standard error. Dashed vertical lines indicate reversal point. Machine reversal curves are not shown.

## Appendix D

### Parameter recovery procedure

We used simulations to verify that the pattern of learning rates obtained in Experiments 1 and 2 did not arise artificially from the parameter optimization procedure. We ran a parameter recovery analysis for discrete sets of parameter values. For Experiments 1 and 2, we simulated 36 virtual participants on our behavioural tasks (36 being the total number of participants in both experiments) with different patterns of learning rates (see **Table S1**). The other parameters were set to their mean fitted values across participants and conditions. The results of these analyses confirmed the capacity of our parameter optimization procedure to correctly recover the true parameters in all experimental conditions.

### Performance of the CF model using different parameter values

We simulated the CF model (see “Modelling” section, main text) in two different environments, where divergence was maximal between buttons (‘environment 1’) or between machines (‘environment 2’) (see **Figure 13A**). We tested the performance of the model with three different patterns of factual and counterfactual alpha rates (‘subjects’, ‘flat’, ‘reverse’). Parameter values were varied according to **Table S1**, below. For the CF simulations (see **Figure 13B)** parameter values from the ‘subjects’ pattern were used, while for the RL simulations parameter values from the CF model were used (α = 0.5, *β* = 0.2, *ρ_machine_* = 8, *ρ_button_ =* 5).

‘Environment 1’ corresponded to the “dependency” condition in Experiment 1, whereas ‘Environment 2’ was similar to the “value” condition in the same experiment. For the simulations, we used the same task structure as the one experienced by the 16 human participants of our sample, but the model generated its own response, and received the outcome corresponding to this response, on each trial. Each model was simulated 10 times, for each environment and pattern.

**Table S1.** Parameter recovery procedure: model parameters used to generate model’s performance, as shown on **Figures 13A**. Parameters of the model are: the (factual) learning rate *α_F_*, the 3 counterfactual learning rates α*_CF_*_1_, α*_CF_*_2_ and α*_CF_*_3_, the reference point *P*, the exploitation intensity *β* and the 2 perseveration biases *ρ_machine_* and *ρ_button_*. Note that only the values of the counterfactual learning rates differed across the ‘subjects’, ‘flat’ and ‘reverse’ simulations.

| Simulations | $\alpha_F$ | $\alpha_{CF1}$ | $\alpha_{CF2}$ | $\alpha_{CF3}$ | $P$ | $\beta$ | $\rho_{machine}$ | $\rho_{button}$ |
| --- | --- | --- | --- | --- | --- | --- | --- | --- |
| ‘subjects’ | 0.5 | 0.25 | 0.5 | 0.5 | 50 | 0.2 | 8 | 5 |
| ‘flat’ | 0.5 | 0.5 | 0.5 | 0.5 | 50 | 0.2 | 8 | 5 |
| ‘reverse’ | 0.5 | 0.5 | 0.25 | 0.25 | 50 | 0.2 | 8 | 5 |

## Footnotes

1 The need to be and feel in control is so strong that individuals do whatever they can to re-establish control when it disappears or is taken away (Brehm, 1966; Brehm and Brehm, 1981). Reestablishment of lost agency can take different forms from illusory pattern perception to erroneous identification of a causal relationship between random or unrelated stimuli. Thus, people experiencing a loss of control are more likely to see images in noise, to form illusory correlations, to perceive conspiracies or to develop superstitions (Whitson & Galinsky, 2008). Such erroneous causal attributions would help restore feelings of control in helplessness individuals by returning the world to a predictable state where “being in control” is the default (Pittman & Pittman, 1980).

2 An irrepressible tendency for playful and exploratory behaviours parallels Hendrick’s “instinct to master”, whose aim is merely “pleasure in exercising a function successfully, regardless of its sensual value”. This “primary pleasure” would arise when efficient action enables the individual to control and alter his environment (Hendrick, 1943). Interestingly, such exploratory behaviours have been found to be more frequent in younger animals, which have less experience with action-outcomes relationships (Siwak, 2001).

3 In a similar vein, it has been suggested that some of our causal beliefs –e.g., control and self-efficacy beliefs– would have evolved to foster the discovery of unpredicted sensory events for which our actions are responsible, which would reinforce and prioritize those actions that lead to control over the environment (Redgrave and Gurney, 2006; Karsh and Eitam, 2015).

4 Here we draw upon a classical distinction between associative and generative approaches to causation, according to which causes are “associated” with effects by retrospection or actively “generate” their effects through an operant mechanism (e.g., Cheng, 1997). Strictly speaking, however, associative models in the form of *reinforcement-learning (RL) algorithms* do also possess a generative model of the world (i.e., an explanation for how observations are generated), whereas *counterfactual emulation* is a generative mechanism per se (i.e., a mechanism to decide which among several candidate causes has generated the effect). Here, and in what follows, we take the “generative” term in a broader and more liberal sense: generative models are those models drawing on an explicit representation of the generative source (usually in the form of a probability distribution over action outcomes), which can be used to make predictions about future outcome states. Representation of the generative source is either complete or approximating the complete solution (that is, an exhaustive representation of all possible action-outcome contingencies). In this sense, generative models are also often normative, i.e., derived from rational principles, and aim at statistical optimality (Gershman, 2015, “normative statistical perspective”).

5 Although the term “simulation” is routinely used to describe the process of running (mental) alternatives to the current situation, the specificity of the counterfactual approach is perhaps best captured by a former distinction between *simulation* and *emulation*, such as examples found in computer science (e.g., Guruprasad, Ricci and Lepreau, 2005). A *simulation* represents a target’s behaviour by explicitly modelling its underlying states, which usually occurs through a generative model known to best represent the actual states at play. Importantly, however, a simulation does not imply to faithfully mimic the outward behaviour of a target (e.g., a simulation may run faster than real time). *Emulation*, conversely, aims to mimic the observable behaviour without having to accurately represent its internal states but ultimately aims to serve as a substitute for the target being emulated (a function - e.g., face recognition - emulated by a neural network). Note that emulating an agent, or a function, is useful when one does not exactly know its internal states, or when representing them accurately would be too demanding. Simulation and emulation hinge on two different assumptions, which align snugly with the generative and counterfactual approaches to causation, respectively. The generative view assumes that individuals infer causation by simulating the internal process through which hidden states generate observable effects. The process is computationally ruinous, but it may provide an accurate estimate on the likelihood of a candidate cause given what is observed. The CF view, on the other hand, does not make any reference to the generative source behind observation. Thus, contrary to generative models that simulate all possible contingencies from a given situation, CF operates by emulating the unchosen alternative only, and by making decisions based on variations of some parameters value (e.g., learning rates) when one travels from the real (factual) to the emulated (counterfactual) world (see Lucas and Kemp, 2015).

